# Gut *Roseburia hominis*-derived 10-hydroxystearic acid alleviates metabolic disorder by restoring hepatic fatty acid metabolism

**DOI:** 10.64898/2026.11.04.764162

**Authors:** Jingxiang Zhang, Chunlei Zhang, Clare Hsu, Minting Hu, Ling Hu, Shaun Tan, Kazuyuki Kasahara, Yifei Ge, Shuli Zhang, Sunny Wong, Rinkoo Dalan, Nguan Soon Tan, Yusuf Ali, Ju-Sheng Zheng, Chuchu Wang, Xiaotao Shen

**Affiliations:** Lee Kong Chian School of Medicine, Nanyang Technological University, Singapore, Singapore; School of Chemistry, Chemical Engineering, and Biotechnology, Nanyang Technological University, Singapore, Singapore; Guizhou Province Livestock and Poultry Genetic Resources Management Station, Guiyang, China; Department of Neurology, The Second Affiliated Hospital of Guangxi Medical University, Nanning, Guangxi, China; KLARIS Precision Health Research Clinic, Department of Endocrinology, Tan Tock Seng Hospital, Singapore, Singapore; School of Biological Sciences, Nanyang Technological University, Singapore, Singapore; School of Life Sciences, Westlake University, Hangzhou, China; NTU Institute of Structural Biology, Nanyang Technological University, Singapore, Singapore; Singapore Phenome Center, Nanyang Technological University, Singapore, Singapore

## Abstract

Metabolic disorders and health-promoting dietary interventions represent complementary contexts of metabolic impairment and improvement, providing an opportunity to identify gut microbial taxa associated with both disease status and dietary response. Here, we integrated five independent human cohorts spanning metabolic disorders and dietary interventions and identified *Roseburia hominis* as a gut microbial taxon that consistently decreased in metabolic disorders and increased after dietary interventions. Beyond species abundance, strain-level genetic variation in *R. hominis* provided additional information for distinguishing metabolic disorder status and stratifying cardiometabolic severity. Integration of phenotype-associated variants with circulating metabolomics prioritized *ohyA*, encoding an oleate hydratase, as a candidate functional determinant linking *R. hominis* genetic variation to host metabolic heterogeneity. An *ohyA*-based genetic burden score was associated with adverse cardiometabolic traits and changed longitudinally during dietary intervention, while circulating metabolites statistically accounted for part of the associations between *ohyA* variants and host phenotypes. Functional experiments demonstrated that *R. hominis* OhyA protein converts oleic acid to 10-hydroxystearic acid (10-HSA), whereas an OhyA carrying five prioritized missense substitutions showed markedly impaired enzymatic activity. In high-fat diet-fed mice, 10-HSA attenuated body-weight gain, improved glucose homeostasis and insulin sensitivity, and reduced dyslipidemia and hepatic steatosis. Multi-omics profiling revealed limited remodeling of the gut microbiota but pronounced hepatic transcriptional and metabolic responses, characterized by activation of PPARα-associated lipid-metabolic pathways, altered fatty-acid oxidation and desaturation, and remodeling of hepatic lipid composition. Together, these findings establish a diet-responsive *R. hominis**–***OhyA***–***10-HSA axis linking microbial variation to host metabolic dysfunction and show that 10-HSA can improve metabolic health in mice.

## Introduction

Metabolic disorders, including prediabetes and type 2 diabetes (T2D), represent a growing global health burden and substantially increase the risk of cardiovascular disease and other long-term complications^1^. This burden is expected to rise further with population aging, urbanization, sedentary lifestyles, and ongoing dietary transitions^2^. Dietary modification is therefore a central component of both the prevention and clinical management of metabolic disorders, and dietary interventions can improve glycemic control, insulin sensitivity, body weight, and broader metabolic health^3^. However, the magnitude and nature of these benefits vary considerably among individuals. Understanding the biological factors that contribute to this heterogeneity is therefore important for elucidating how diet influences metabolic health and for developing more effective dietary strategies.

The gut microbiota has emerged as an important interface between dietary exposures and host metabolism. Its composition and function are strongly shaped by diet, medication use, and other environmental factors^4–6^, while gut microbes can in turn transform dietary components and generate bioactive metabolites that influence host physiology^7–10^. Perturbations of the gut microbiome have been associated with a broad range of metabolic diseases, including obesity, T2D, and metabolic dysfunction-associated steatotic liver disease^11,12^. However, disease-associated and diet-responsive microbiome changes are commonly investigated separately, and both show substantial heterogeneity across individuals and populations. Integrating these two contexts may provide a more stringent strategy for identifying microbial taxa relevant to metabolic health: taxa that are reproducibly altered during metabolic impairment and shift in the opposite direction after dietary intervention may represent particularly informative components of the diet–microbiome–host metabolic interface.

Integrating disease-associated and diet-responsive changes can help identify microbial taxa that are particularly relevant to metabolic health. Beyond these species-level abundance patterns, considerable genetic diversity exists within microbial species^13^, and closely related strains can differ in gene content, copy number, genome organization, nucleotide sequence, and single-nucleotide polymorphisms (SNPs)^14–17^. Such variation can modify microbial proteins and metabolic pathways, potentially altering substrate utilization and metabolite production even in the absence of detectable differences in species abundance. Consistent with this concept, *Prevotella copri* comprises genetically distinct clades with pronounced geographical and evolutionary structure^18^, whereas disease-associated genomic differences have been observed among *Ruminococcus gnavus* isolates from individuals with Crohn’s disease^19,20^. Microbial SNPs have also been associated with host characteristics including body mass index (BMI), dietary patterns, and infection status^21–23^. Thus, species abundance and strain-level genetic variation may provide complementary information about microbial function: abundance reflects the representation of a microbial population within the community, whereas genetic variation may further shape its biochemical capacity. Despite these advances, a major gap remains between strain-resolved association studies and mechanistic understanding. It is still poorly understood whether microbial genetic variation alters the activity of specific metabolic enzymes, whether these changes modify production of host-accessible metabolites, and whether such functional variation contributes to inter-individual differences in metabolic health. More broadly, few studies have established a continuous experimental link from diet-responsive microbial ecology to microbial genetic variation, microbial enzyme activity, metabolite production, and downstream host metabolic physiology.

Here, we addressed these questions through an integrative discovery and experimental framework spanning human cohorts, microbial genomics, metabolomics, biochemical validation, and *in vivo* functional studies. We first combined five independent human cohorts, including two metabolic disorder cohorts^24,25^ and three dietary intervention cohorts^7,8^, to identify gut microbial taxa showing reciprocal changes across metabolic deterioration and dietary improvement. This multi-cohort strategy prioritized *Roseburia hominis*, which consistently decreased in individuals with metabolic disorders and increased after dietary intervention. We then examined whether variation within *R. hominis* itself contributed additional functional information beyond species abundance. Integrating strain-level genetic variation with host cardiometabolic traits and circulating metabolomics revealed substantial associations between *R. hominis* genetic variation and host metabolic heterogeneity and prioritized the *ohyA* locus as a candidate functional determinant. We further showed that *ohyA* encodes an oleate hydratase that converts oleic acid (OA) to 10-hydroxystearic acid (10-HSA), whereas an OhyA variant carrying five prioritized amino acid substitutions exhibited markedly impaired enzymatic activity. To determine whether this microbial metabolic product contributes to host metabolic regulation, we administered 10-HSA to high-fat diet-fed mice and found that it ameliorated multiple features of high-fat diet (HFD)-induced metabolic dysfunction. Finally, integrated transcriptomic, protein, and metabolomic profiling indicated that the metabolic effects of 10-HSA were accompanied by pronounced hepatic lipid remodeling and activation of a PPARα-associated metabolic program. Together, these findings define a diet-responsive *R. hominis**–***OhyA***–***10-HSA axis that links microbial abundance and strain-level functional variation to microbial lipid metabolism and host metabolic health.

## Results

### Multi-cohort analysis identifies *R. hominis* as a diet-responsive gut microbial taxon associated with metabolic health

To identify gut microbial taxa robustly associated with metabolic disorders, we designed an integrative multi-cohort study combining metabolic disorder cohorts^24,25^ with dietary intervention cohorts^7,8^ (Fig. 1a). We reasoned that microbial taxa consistently altered in metabolic disorders and modulated in the opposite direction by dietary intervention would represent strong candidates linking the gut microbiome to metabolic health.

**Figure 1:**
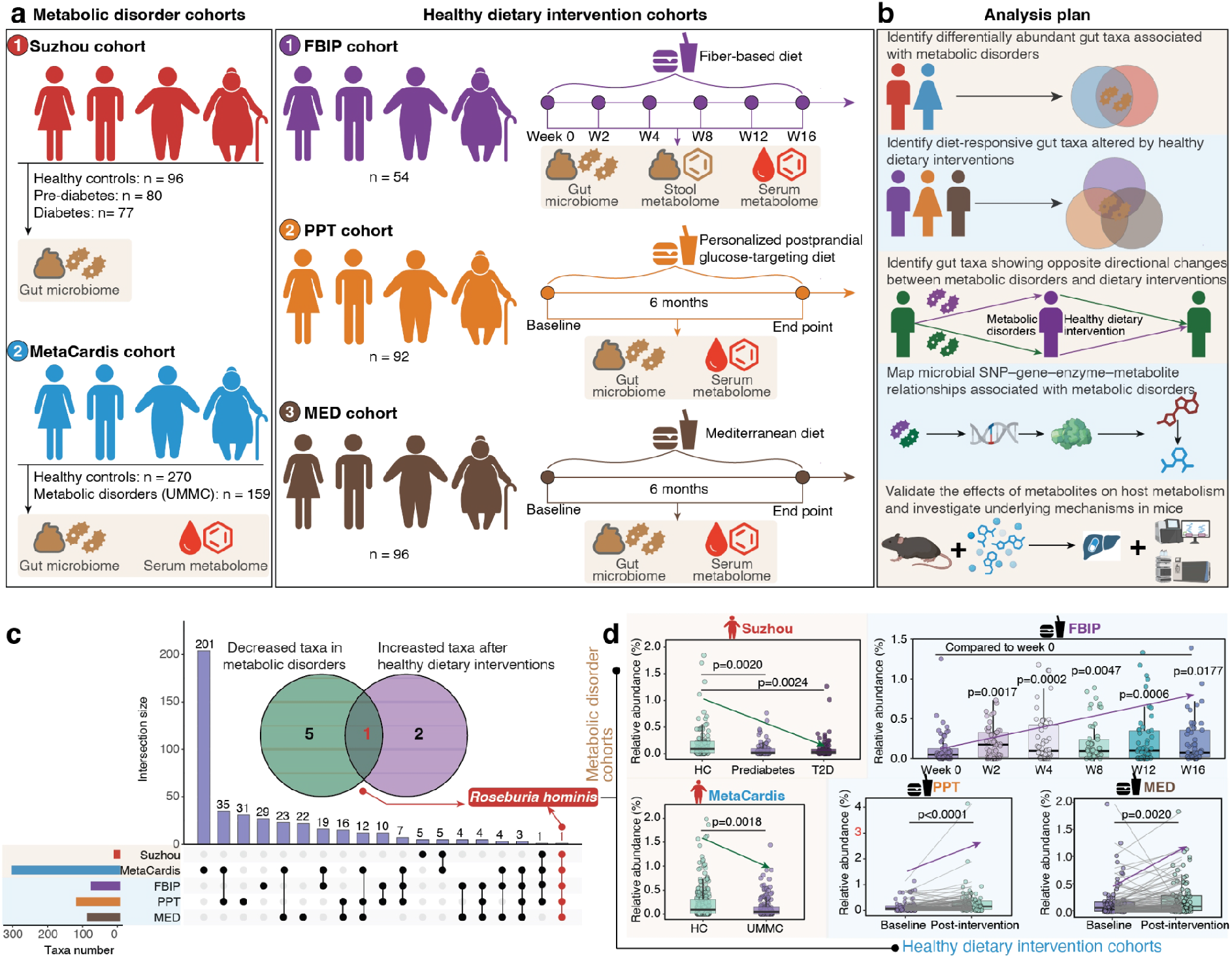
Integrative multi-cohort analysis identifies *R. hominis* as a diet-responsive gut microbial taxon associated with metabolic health. (a) Overview of the five independent human cohorts included in the study, comprising two metabolic disorder cohorts (Suzhou and MetaCardis) and three dietary intervention cohorts: the fiber-based intervention program (FBIP), personalized postprandial glucose-targeting diet (PPT), and Mediterranean diet (MED) cohorts. (b) Schematic overview of the integrative analytical framework used to identify gut microbial taxa consistently associated with metabolic disorders and showing reciprocal changes after dietary intervention, followed by strain-level genetic, metabolic, and experimental analyses. (c) Intersection of gut microbial taxa consistently decreased in the two metabolic disorder cohorts and taxa consistently increased after dietary intervention across the three cohorts. *R. hominis* was the only taxon meeting both criteria across all five cohorts. (d) Relative abundance of *R. hominis* across metabolic disorder groups in the Suzhou and MetaCardis cohorts (left) and before and after dietary intervention in the FBIP, PPT and MED cohorts (right).

We assembled five independent human cohorts comprising 924 participants in total (Fig. 1a, Supplementary Fig. 1a and Supplementary Table 1; Methods). Two metabolic disorder cohorts were included: the Suzhou cohort^24,25^ (n = 253), comprising healthy controls (n = 96) and individuals with prediabetes (n = 80) or type 2 diabetes (T2D, n = 77), and the MetaCardis cohort^24,25^ (n = 429), comprising healthy controls (n = 270) and untreated metabolically matched controls (UMMC, n = 159), who had metabolic syndrome but were not receiving relevant medication (Methods). We additionally included three dietary intervention cohorts: fiber-based intervention program (FBIP, n = 54)^7^, personalized postprandial glucose-targeting diet (PPT, n = 92)^8^, and Mediterranean diet (MED, n = 96).

Based on this integrative design, we established a stepwise analytical framework to identify microbial taxa showing concordant changes across disease cohorts and reciprocal responses across dietary intervention cohorts (Fig. 1b). We first performed differential abundance analyses in the two metabolic disorder cohorts (Supplementary Fig. 1b and Methods). We identified six species-level gut microbial taxa that were consistently decreased in individuals with metabolic disorders in both cohorts (Supplementary Fig. 1c, d). These included *GGB9677_SGB15180*, *Roseburia hominis*, *Lawsonibacter hominis*, *Blautia faecicola*, *Clostridium sp. AM33_3* and *GGB51512_SGB71885* (Supplementary Fig. 1d).

We next investigated microbial taxa responsive to dietary interventions by analyzing three dietary intervention cohorts (Methods). This analysis revealed three microbial taxa that were consistently increased after dietary intervention, including *Clostridium_SGB4714*, *Butyribacter intestini,* and *R. hominis* (Supplementary Fig. 1e, f). We then intersected metabolic disorder-associated taxa with diet-responsive taxa to identify microbial taxa showing opposite directional changes across disease and intervention contexts. Strikingly, only a single microbial taxon, *R. hominis*, met both criteria across all five cohorts analyzed (Fig. 1c). Notably, *R. hominis* was consistently decreased in metabolic disorder cohorts, while showing a reproducible increase after dietary interventions (Fig. 1d). Together, these analyses identify *R. hominis* as a robust and diet-responsive microbial taxon associated with host metabolic health and motivate subsequent strain-level genetic and functional analyses.

### Cardiometabolic trait-associated genetic variation in *R. hominis* distinguishes metabolic disorder status

To further investigate whether strain-level genetic variation in *R. hominis* contributes to its association with host metabolic health, we next focused on the MetaCardis cohort, which provided both metagenomic data and detailed cardiometabolic phenotyping (Methods). We performed SNP calling for *R. hominis*, identifying 97,252 SNPs that were subsequently tested for associations with eight cardiometabolic traits, spanning adiposity (BMI and waist circumference), glycemic control (HbA1c), lipid metabolism (triglycerides), systemic inflammation (C-reactive protein [CRP]), adipokine status (adiponectin), and blood pressure (systolic and diastolic blood pressure [SBP and DBP]) (Methods). Across the eight cardiometabolic traits, 5,715 distinct SNPs ranked within the top 1% of association signals for at least one host trait (Supplementary Fig. 2a, b and Methods). To identify genes showing recurrent genetic associations across cardiometabolic traits, we retained genes harboring top-ranked SNP association signals for seven or all eight traits. Applying this criterion identified 603 distinct SNPs distributed across 68 genes (Fig. 2a and Supplementary Table 2), indicating that a subset of *R. hominis* genetic variants was consistently associated with multiple dimensions of host cardiometabolic status.

**Figure 2:**
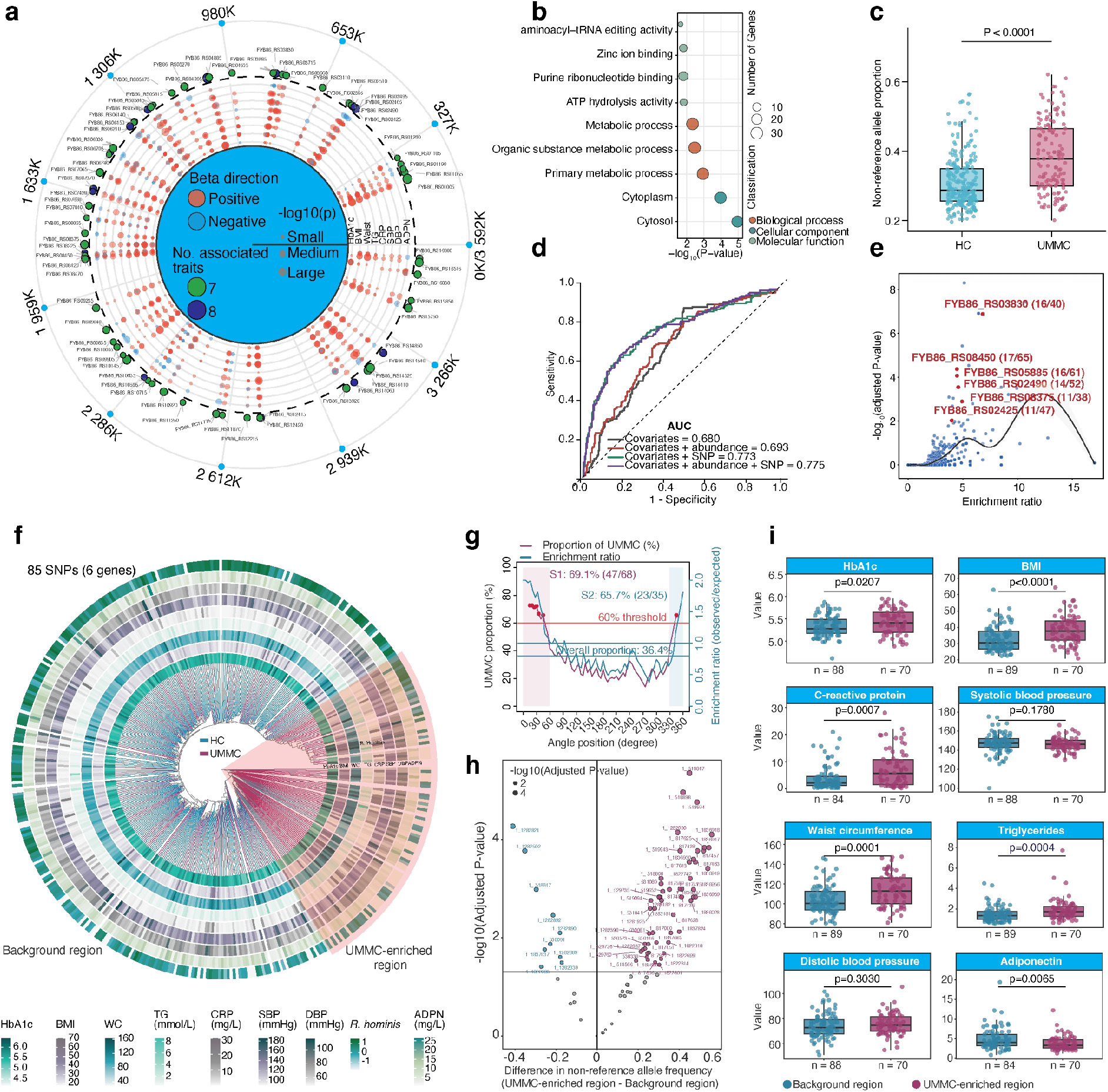
Cardiometabolic trait-associated SNPs in *R. hominis* distinguish metabolic disorder status and stratify disease severity. (a) Associations between *R. hominis* SNPs and eight host cardiometabolic traits. SNPs ranked within the top 1% of association signals for at least seven of the eight cardiometabolic traits are highlighted. (b) Functional enrichment analysis of genes harboring the cardiometabolic trait-associated SNPs identified in (a). (c) Proportion of non-reference alleles in *R. hominis* in healthy controls (HC) and participants with metabolic disorders (UMMC). (d) ROC curves for discrimination of metabolic disorder status using models incorporating host covariates alone, host covariates plus *R. hominis* relative abundance, host covariates plus *R. hominis* SNPs, or host covariates plus both abundance and SNPs. (e) Enrichment analysis identifying six *R. hominis* genes disproportionately harboring SNPs associated with host cardiometabolic traits. (f) Sample-level phylogenetic tree reconstructed using 85 candidate SNPs located within the six genes identified in (e). (g) Sliding-window analysis of the proportion of UMMC across the tree and the corresponding observed-to-expected enrichment ratio. (h) Differences in non-reference allele frequencies of the 85 candidate SNPs between UMMC participants in UMMC-enriched and background regions. (i) Comparison of eight cardiometabolic traits between UMMC participants in the UMMC-enriched and background regions. Six of the eight traits differed significantly between the two groups.

To explore the potential functional relevance of these variants, we performed enrichment analysis of the genes harboring the cardiometabolic trait-associated SNPs^26^ (Methods). The enriched annotations were primarily related to metabolic functions, including metabolic process, organic substance metabolic process, and primary metabolic process (Fig. 2b; FDR-adjusted *P* < 0.05), suggesting that genes harboring broadly trait-associated SNPs were related to metabolic processes. We next examined whether the overall non-reference allele proportion differed between healthy controls and participants with metabolic disorders. Notably, participants with metabolic disorders exhibited a significantly higher proportion of non-reference alleles than healthy controls (Fig. 2c; Wilcoxon rank-sum test, *P* < 0.0001).

To quantitatively assess whether within-species genetic variation provided additional information beyond species-level relative abundance, we constructed random forest classification models incorporating host covariates (country, sex, and age) alone, or in combination with *R. hominis* relative abundance, SNP profiles, or both (Methods). The covariate-only model achieved an AUC of 0.680, which increased modestly to 0.693 after inclusion of *R. hominis* relative abundance. Incorporation of the 603-SNP profile resulted in a higher AUC of 0.773, while further addition of *R. hominis* relative abundance yielded a similar AUC of 0.775 (Fig. 2d). Furthermore, PCA based on *R. hominis* SNP profiles revealed separation between UMMC participants and healthy controls, as supported by PERMANOVA (*P* = 0.001, R^2^ = 0.020; Supplementary Fig. 2c, d; Methods).

Together, these results demonstrate that strain-level genetic variation in *R. hominis* is associated with host cardiometabolic traits and provides additional information on metabolic disorder status beyond species-level relative abundance.

### SNP-defined *R. hominis* genetic profiles stratify cardiometabolic heterogeneity

We next sought to determine whether a subset of cardiometabolic trait-associated *R. hominis* variants could capture metabolic heterogeneity among participants with metabolic disorders. We therefore prioritized genes disproportionately enriched for SNPs associated with host cardiometabolic traits using a hypergeometric enrichment test (Methods). Six genes, *FYB86_RS03830*, *FYB86_RS08450*, *FYB86_RS05885*, *FYB86_RS02490*, *FYB86_RS08375*, and *FYB86_RS02425*, showed significant enrichment and collectively contained 85 candidate SNPs associated with all eight cardiometabolic traits (Fig. 2e). These analyses defined a focused *R. hominis* genetic signature for subsequent participant-level stratification.

We then reconstructed a sample-level phylogenetic tree using the 85 SNPs located within these 6 genes (Methods). The resulting topology showed partial segregation of UMMC participants and healthy controls based solely on their *R. hominis* SNP profiles (Fig. 2f). Notably, sliding-window enrichment analysis identified two contiguous segments enriched for UMMC participants, in which UMMC participants accounted for 69.1% (47/68) and 65.7% (23/35) of participants, respectively, compared with an overall proportion of 36.4% across the tree (Fig. 2g and Methods). We collectively designated these two UMMC-enriched segments as the UMMC-enriched region and the remainder of the tree as the background region (Fig. 2f).

To determine whether UMMC participants in the UMMC-enriched and background regions harbored distinct *R. hominis* genetic profiles, we compared non-reference allele frequencies of the 85 candidate SNPs between the two groups (Methods). Most SNPs (63/85) differed significantly between the groups, with the majority of these differential variants (53/63) showing higher non-reference allele frequencies in the UMMC-enriched region (Fisher’s exact test with 10,000 Monte Carlo simulations, FDR-adjusted *P* < 0.05; Fig. 2h). These findings indicate that the two UMMC subgroups are characterized by distinct patterns of *R. hominis* genetic variation. Notably, the three SNPs showing the strongest statistical differences, defined as those with the three lowest FDR-adjusted *P* values among the 85 candidate SNPs, were located at positions 519847, 518898, and 519974, all within the *FYB86_RS02425* gene (Fig. 2h).

We next asked whether these distinct *R. hominis* genetic profiles were accompanied by differences in host cardiometabolic status. UMMC participants in the UMMC-enriched region exhibited higher HbA1c, BMI, CRP, waist circumference, and triglyceride levels, together with lower adiponectin levels, than those in the background region, whereas systolic and diastolic blood pressure did not differ significantly between the groups (Fig. 2i). The concordant differences across six of the eight cardiometabolic traits indicate a more adverse cardiometabolic profile among UMMC participants in the UMMC-enriched region, consistent with greater metabolic disorder severity.

Together, these findings demonstrate that SNP-defined *R. hominis* genetic subpopulations are associated with distinct cardiometabolic profiles among participants with metabolic disorders, linking strain-level microbial genetic heterogeneity to metabolic disorder severity.

### *ohyA* emerges as a key *R. hominis* gene linking microbial genetic variation to host metabolomic variation

We next investigated whether cardiometabolic trait-associated *R. hominis* variants were also associated with circulating metabolomic profiles. Because microbial genetic variation can influence bacterial metabolic activity and downstream microbial metabolic outputs^27^, we hypothesized that integrating *R. hominis* genetic variation with circulating metabolomics could help prioritize microbial genes potentially linked to host metabolic phenotypes^28^.

We first examined associations between the 85 SNPs identified above and circulating metabolites in the MetaCardis cohort (Methods). This analysis revealed numerous SNP-metabolite associations involving diverse metabolite classes, including bile acids, amino acids, aromatic microbial metabolites, and host-microbe co-metabolites (Fig. 3a). These findings indicate that strain-level genetic variation in *R. hominis* is associated with variation across multiple classes of circulating metabolites.

**Figure 3:**
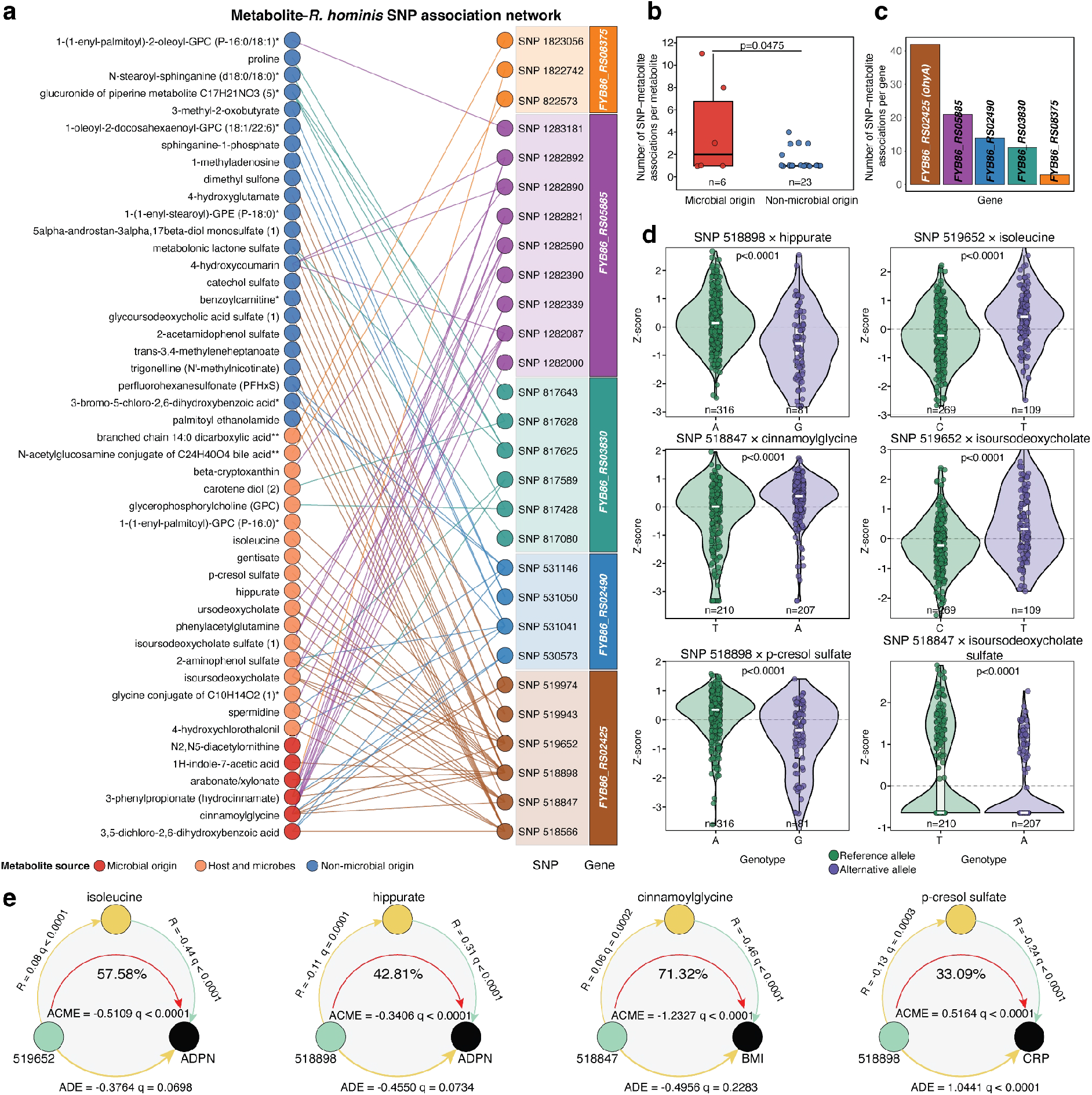
*ohyA* emerges as a key *R. hominis* gene linking microbial genetic variation to host metabolomic variation. (a) Association network between the 85 candidate *R. hominis* SNPs and circulating metabolites. (b) Number of SNP-metabolite associations per metabolite for microbial-origin and non-microbial-origin metabolites. (c) Number of SNP-metabolite associations attributable to each SNP-containing gene. *FYB86_RS02425* (*ohyA*) exhibited the largest number of SNP-metabolite associations. (d) Representative circulating metabolite levels stratified by *R. hominis* SNP alleles. (e) Statistical mediation analyses of selected *ohyA* SNP-metabolite-cardiometabolic trait relationships.

We next asked whether these associations preferentially involved metabolites with potential microbial contributions. Metabolites classified as microbial-origin exhibited a greater number of SNP-metabolite associations per metabolite than non-microbial-origin metabolites (Fig. 3b; Wilcoxon rank-sum test, *P* = 0.0475), suggesting preferential involvement of metabolites with predicted microbial contributions. We then quantified the number of SNP-metabolite associations attributable to each of the six candidate genes identified in the preceding analysis. Among these genes, *FYB86_RS02425* (*ohyA* [oleate hydratase]) showed the largest number of SNP-metabolite associations (n = 42; Fig. 3c). Notably, *ohyA* also harbored several SNPs showing pronounced allele-frequency differences between UMMC participants in the UMMC-enriched and background regions (Fig. 2h). Thus, complementary analyses of host cardiometabolic traits, circulating metabolites, and disease-associated SNP patterns converged on *ohyA* as a leading candidate gene linking *R. hominis* genetic variation to host metabolism.

We next analyzed representative *ohyA* SNP-metabolite associations at the genotype level (Methods). SNP 518898 was associated with differences in circulating hippurate and p-cresol sulfate levels, whereas SNP 519652 was associated with isoleucine and isoursodeoxycholate levels (Fig. 3d). Similarly, SNP 518847 was associated with cinnamoylglycine and isoursodeoxycholate sulfate levels (Fig. 3d). We then investigated whether metabolites associated with these *ohyA* variants were also related to the corresponding host cardiometabolic traits. Hippurate was positively associated with adiponectin, whereas isoleucine was negatively associated with adiponectin. Cinnamoylglycine was negatively associated with BMI, and p-cresol sulfate was negatively associated with CRP (Supplementary Fig. 3). Together, these analyses identified a set of SNP-metabolite-trait relationships connecting *ohyA* variation with host cardiometabolic traits.

Furthermore, we used mediation analysis to assess whether these circulating metabolites could statistically account for part of the associations between *ohyA* variants and host cardiometabolic traits (Methods). Significant indirect effects were observed for all four tested SNP-metabolite-trait relationships (Fig. 3e). Isoleucine statistically accounted for an estimated 57.6% of the association between SNP 519652 and adiponectin (ACME = -0.5109, FDR-adjusted *P* < 0.0001), whereas hippurate accounted for 42.8% of the association between SNP 518898 and adiponectin (ACME = - 0.3406, FDR-adjusted *P* < 0.0001). Cinnamoylglycine accounted for 71.3% of the association between SNP 518847 and BMI (ACME = -1.2327, FDR-adjusted *P* < 0.0001), and p-cresol sulfate accounted for 33.1% of the association between SNP 518898 and CRP (ACME = 0.5164, FDR-adjusted *P* < 0.0001).

Together, these findings prioritize *ohyA* as a key *R. hominis* candidate gene at the intersection of microbial genetic variation, circulating metabolomic variation, and host cardiometabolic traits. The observed SNP-metabolite-trait relationships are consistent with a statistical model in which circulating metabolites form an intermediate molecular layer between *ohyA* variation and host cardiometabolic heterogeneity.

### *ohyA* gene-level SNP burden is associated with adverse cardiometabolic traits and responds to dietary intervention

To further investigate whether the aggregate genetic variation within the *ohyA* locus was associated with host metabolic status and responsiveness to dietary intervention, we developed a derived genetic burden score (dGBS). Because individual SNPs capture only a fraction of the genetic variation within a gene, the dGBS integrates the cumulative contribution of multiple *ohyA* variants at the individual level (Fig. 4a and Methods).

**Figure 4:**
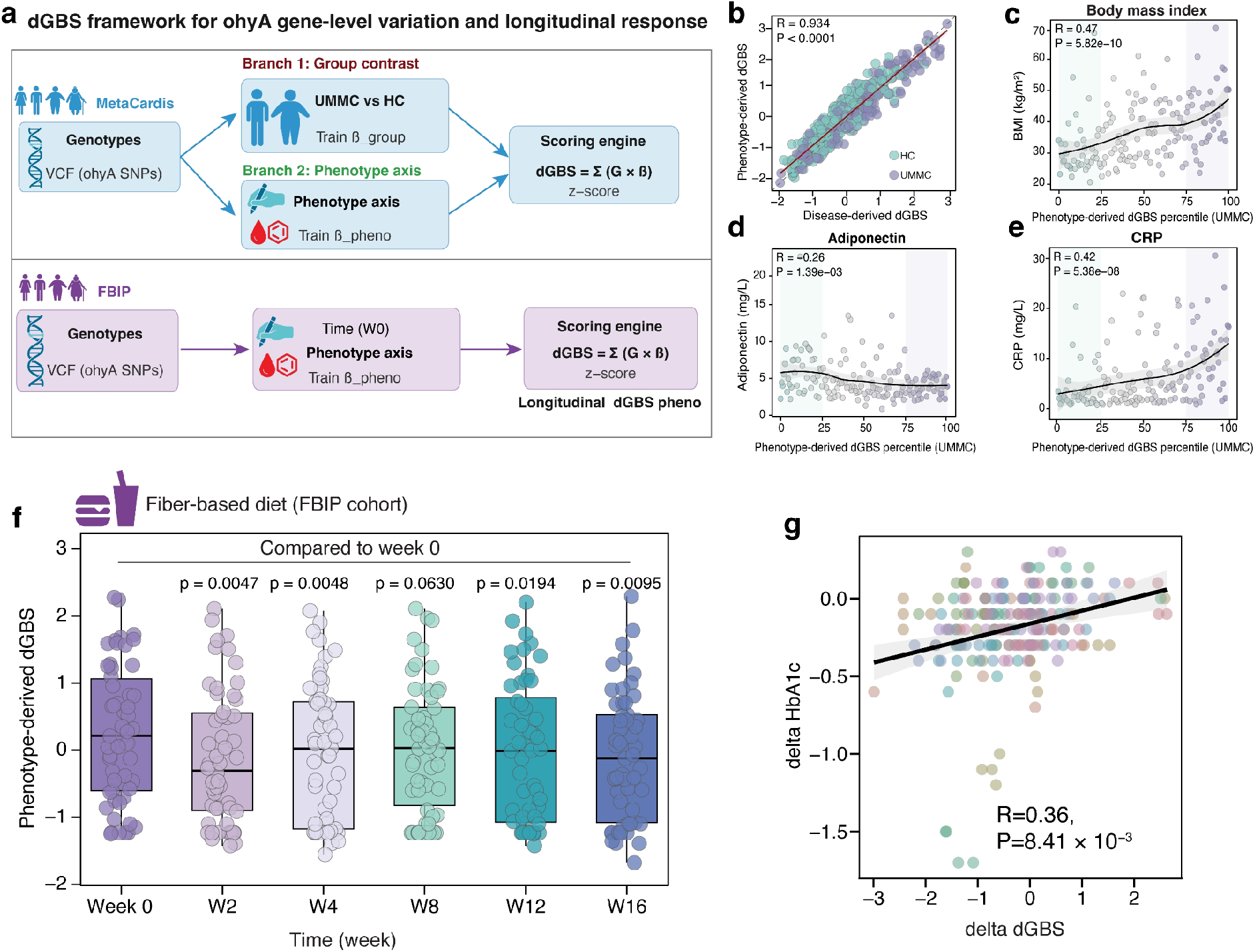
*ohyA* gene-level SNP burden in *R. hominis* is associated with host metabolic risk and dietary response. (a) Framework for constructing the *ohyA* gene-level derived genetic burden score (dGBS). In the MetaCardis cohort, disease-derived and phenotype-derived dGBS models were constructed using metabolic disorder status and the cardiometabolic traits axis, respectively. In the longitudinal FBIP cohort, the phenotype-derived dGBS was applied to assess changes in the *ohyA* genetic profile during dietary intervention. (b) Correlation between disease-derived and phenotype-derived dGBS in the MetaCardis cohort. (c-e) Associations of phenotype-derived dGBS with BMI, adiponectin, and CRP among UMMC participants. (f) Longitudinal changes in phenotype-derived *ohyA* dGBS during the dietary intervention in the FBIP cohort. (g) Association between within-participant changes in *ohyA* dGBS and HbA1c during dietary intervention.

In the MetaCardis cohort, we constructed two complementary dGBS models. A disease-derived dGBS was trained using metabolic disorder status as the outcome, whereas a phenotype-derived dGBS was trained using quantitative cardiometabolic traits as outcomes (Methods). These complementary approaches allowed us to assess whether aggregate *ohyA* genetic variation consistently captured both categorical disease status and continuous cardiometabolic variation. The two independently derived scores were strongly correlated (Pearson’s r = 0.934, *P* < 0.0001; Fig. 4b), indicating that they captured highly concordant *ohyA* genetic signatures. Both dGBS measures were significantly higher in UMMC participants than in healthy controls (Supplementary Fig. 4a). Given our previous observation that *ohyA*-associated SNP profiles captured cardiometabolic heterogeneity among UMMC participants (Fig. 2i), we used the phenotype-derived dGBS for subsequent analyses. Within the UMMC group, higher phenotype-derived dGBS values were associated with less favorable cardiometabolic traits. Specifically, participants in the highest dGBS quartile had higher BMI and CRP levels and lower adiponectin concentrations than those in the lowest quartile (Fig. 4c-e). These findings indicate that the aggregate *ohyA* SNP burden captures graded variation in cardiometabolic status beyond that represented by individual variants.

We next applied the dGBS framework to the longitudinal FBIP cohort to determine whether the *ohyA* genetic signature changed during dietary intervention (Fig. 4a). The phenotype-derived dGBS decreased significantly over the course of the intervention (Fig. 4f), indicating a longitudinal shift in the detectable *ohyA* allelic profile toward the pattern associated with a more favorable cardiometabolic state in the MetaCardis cohort. Moreover, within-participant changes in dGBS were significantly associated with changes in HbA1c, with greater reductions in dGBS accompanying greater improvements in glycemic control (Fig. 4g). Changes in dGBS were also associated, although more modestly, with changes in BMI and triglyceride levels (Supplementary Fig. 4b). Thus, the *ohyA* allele profile was associated with diet-related metabolic changes during the intervention. These longitudinal observations suggest that the within-species genetic composition of *R. hominis* at the *ohyA* locus may vary in association with dietary modulation.

### OhyA functions as an oleate hydratase that converts oleic acid into 10-hydroxystearic acid

o*hyA* emerged as a strong candidate gene linking *R. hominis* genetic variation to host metabolic health; we therefore investigated the functional consequences of all *ohyA* variants. Sixty-four SNPs were identified within the *ohyA* locus, including 49 synonymous and 15 missense variants (Supplementary Fig. 4c, d; Methods). Although synonymous variants may affect gene expression or transcript processing, missense variants are more directly capable of altering protein structure and function^29^. We then evaluated the potential effects of the 15 missense variants using two complementary computational approaches: DDMuT^30^, which predicts the effects of amino acid substitutions on protein stability, and PROVEAN^31^, which estimates their potential functional impact (Methods). Five missense variants, F156L, R458S, V486M, R543S, and R565S, were consistently predicted by both approaches to have potentially deleterious effects on OhyA stability or function (Fig. 5a). Mapping these residues onto the predicted OhyA protein structure (AlphaFold ID: AF-G2T1V8-F1) showed that they were distributed across the protein rather than concentrated within a single structural region (Fig. 5b and Methods).

**Figure 5:**
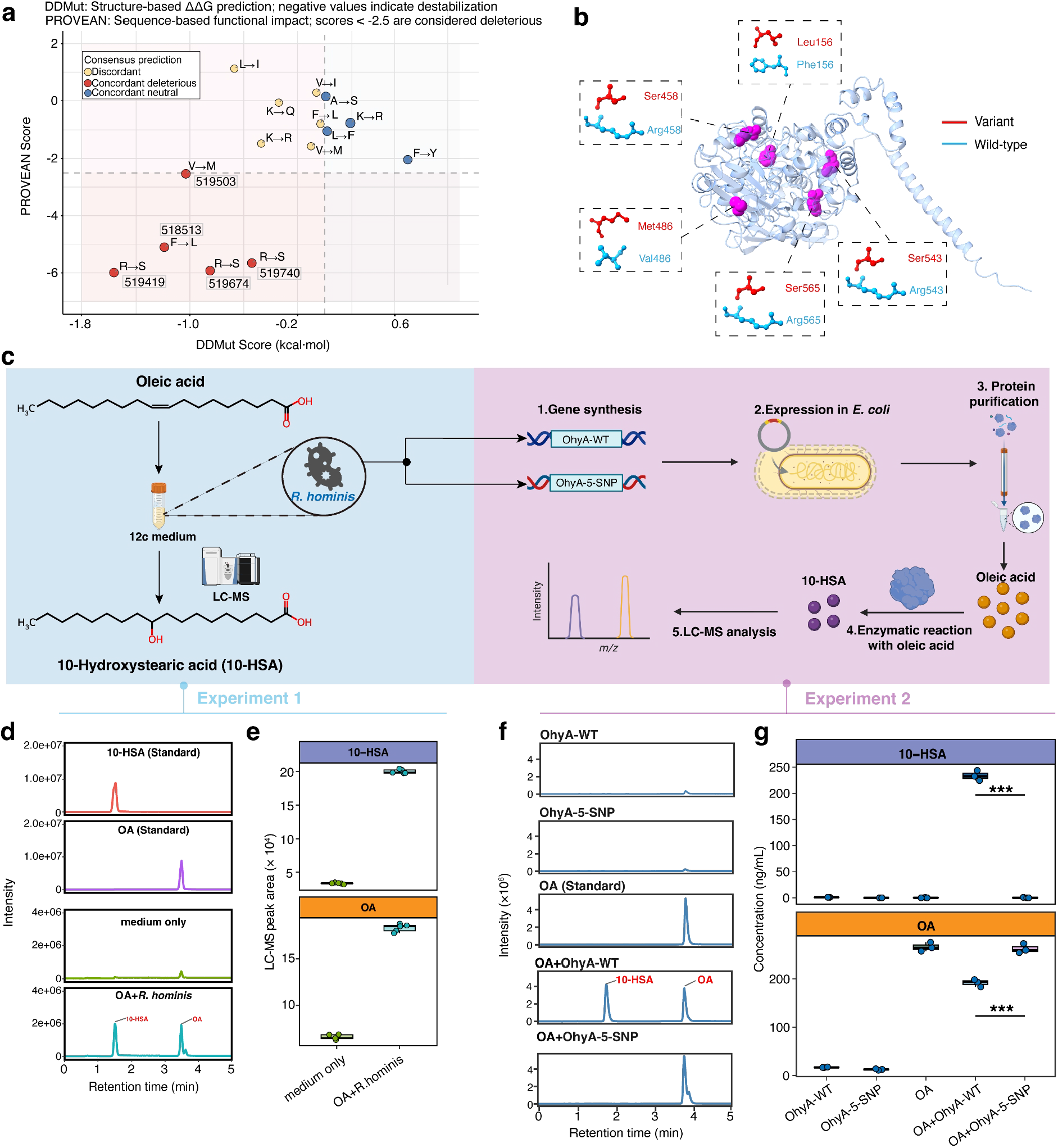
*R. hominis* OhyA converts oleic acid to 10-hydroxystearic acid, whereas the 5-SNP variant shows impaired enzymatic activity. (a) Computational assessment of the 15 missense variants using DDMut and PROVEAN. (b) Structural mapping of the five candidate deleterious variants onto the predicted three-dimensional structure of the OhyA protein. (c) Experimental design for validating the conversion of OA to 10-HSA by *R. hominis* and recombinant OhyA proteins. (d) Representative LC-MS chromatograms showing 10-HSA production following incubation of *R. hominis* with OA, together with authentic OA and 10-HSA standards. (e) Quantification of 10-HSA and residual OA following incubation of *R. hominis* with OA. (f) Representative LC-MS chromatograms showing OA-to-10-HSA conversion in cell-free enzymatic assays with wild-type OhyA (OhyA-WT) or the 5-SNP OhyA variant (OhyA-5-SNP). (g) Quantification of 10-HSA production and residual OA in enzymatic assays containing OhyA-WT or OhyA-5-SNP.

Functional annotation predicted that the *R. hominis ohyA* gene encodes an oleate hydratase that catalyzes the hydration of unsaturated fatty acids [GO:0050151]. We therefore first tested whether *R. hominis* could convert oleate (OA) to 10-hydroxystearic acid (10-HSA) *in vitro*. *R. hominis* was cultured in the presence of OA, and metabolites in the culture supernatant were analyzed by LC-MS (Fig. 5c, left and Methods). Notably, 10-HSA was detected in cultures containing both *R. hominis* and OA but not in bacteria-free medium supplemented with OA (Fig. 5d). Quantitative analysis further showed that incubation with *R. hominis* resulted in a marked increase in 10-HSA accompanied by depletion of OA (Fig. 5e). These results establish that *R. hominis* can convert OA to 10-HSA under the tested culture conditions.

The biochemical activity of the *R. hominis* OhyA protein has not been well characterized. We then asked whether this conversion was directly catalyzed by the OhyA protein and whether the five prioritized variants altered its enzymatic activity (Fig. 5c, right). The *R. hominis ohyA* gene was synthesized and heterologously expressed in *Escherichia coli*, and the purified OhyA protein was incubated with OA in a cell-free enzymatic assay (Supplementary Fig. 5a and Methods). Wild-type OhyA efficiently converted OA to 10-HSA, accompanied by a corresponding reduction in residual OA levels, confirming that *R. hominis* OhyA functions as an oleate hydratase (Fig. 5f, g and Supplementary Fig. 5b). Furthermore, we generated a plasmid encoding an OhyA carrying all top five SNP variants (OhyA-5-SNP). Remarkably, purified OhyA-5-SNP produced substantially less 10-HSA than the wild-type enzyme (Fig. 5f, g).

Together, these experiments establish OhyA as a functional oleate hydratase in *R. hominis* and demonstrate that the disease-associated mutant markedly impairs its ability to convert OA to 10-HSA. These findings provide direct functional support for a model in which strain-level *ohyA* variation alters 10-HSA production and may thereby influence host metabolic health.

### 10-HSA alleviates HFD-induced metabolic dysfunction in mice

We next investigated whether 10-HSA, the metabolic product of *R. hominis* OhyA, could contribute to the beneficial metabolic effects associated with *R. hominis*. To test the effects of 10-HSA *in vivo*, we conducted an 8-week 10-HSA supplementation experiment in C57BL/6 mice (Fig. 6a and Methods). Mice were randomly assigned to four groups receiving a normal diet (ND), ND supplemented with 10-HSA (ND + 10-HSA), a high-fat diet (HFD), or HFD supplemented with 10-HSA (HFD + 10-HSA).

**Figure 6:**
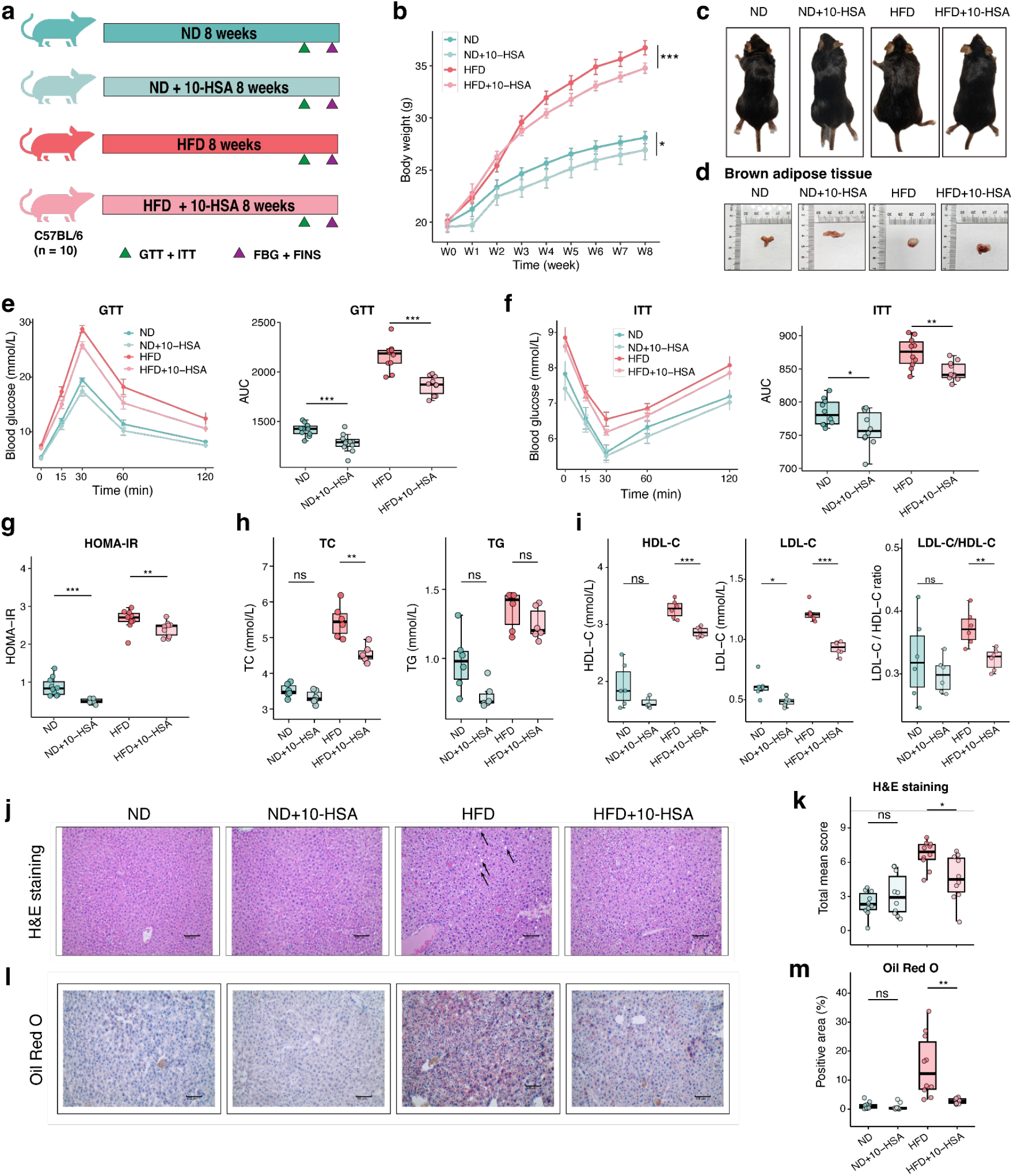
10-HSA alleviates HFD-induced metabolic dysfunction in mice. (a) Experimental design of the 8-week 10-HSA supplementation experiment in mice. (b) Body-weight trajectories during the 8-week intervention. (c) Representative images of mice from the four experimental groups at the study endpoint. (d) Representative images of brown adipose tissue collected at the study endpoint. (e) Glucose tolerance test (GTT) curves and corresponding AUC. (f) Insulin tolerance test (ITT) curves and corresponding AUC. (g) Homeostatic model assessment of insulin resistance (HOMA-IR) across the four groups. (h) Plasma total cholesterol (TC) and triglyceride (TG) levels. (i) Plasma high-density lipoprotein cholesterol (HDL-C) level, low-density lipoprotein cholesterol (LDL-C) level, and LDL-C/HDL-C ratio. (j) Representative H&E-stained liver sections from the four groups. (k) Quantification of histopathological changes in H&E-stained liver sections. (l) Representative Oil Red O-stained liver sections showing hepatic lipid accumulation. (m) Quantification of the Oil Red O-positive area in liver sections.

Within the observation window, HFD feeding induced progressive body-weight gain compared with ND-fed controls. 10-HSA supplementation significantly altered body-weight trajectories in a diet-dependent manner, attenuating HFD-induced weight gain from week 3 onward and resulting in a 1.96 g reduction in final body weight relative to HFD controls; a smaller absolute reduction of 1.18 g was observed under ND conditions (baseline-adjusted linear mixed-effects model,; within-diet treatment × time interactions: ND, *P* = 0.026; HFD, *P* < 0.001; Holm-adjusted post hoc comparisons; Fig. 6b, c). Food intake was monitored throughout the intervention and did not differ significantly between HFD-fed mice with and without 10-HSA supplementation, indicating that the lower body-weight gain associated with 10-HSA treatment was unlikely to be explained by reduced food intake (Supplementary Table 3). Examination of adipose and hepatic tissues further revealed differences between HFD-fed mice with and without 10-HSA supplementation, including an improved brown adipose tissue phenotype in the HFD + 10-HSA group (Fig. 6d and Supplementary Fig. 6a).

We next checked glucose homeostasis and insulin sensitivity. Glucose-tolerance testing showed improved glucose clearance in HFD-fed mice receiving 10-HSA compared with HFD controls (Fig. 6e). Similarly, insulin-tolerance testing indicated improved insulin sensitivity following 10-HSA supplementation (Fig. 6f). Consistent with these dynamic measurements, fasting blood glucose, fasting insulin, and homeostatic model assessment of insulin resistance (HOMA-IR) were significantly lower in HFD + 10-HSA mice than in HFD controls (Welch’s t test; Fig. 6g and Supplementary Fig. 6b, c). Together, these findings indicate that 10-HSA ameliorates HFD-induced disturbances in glucose homeostasis and insulin sensitivity.

We subsequently checked another two metabolic phenotypes: circulating markers of hepatic injury and lipid metabolism. Serum alanine aminotransferase (ALT) and aspartate aminotransferase (AST) levels were lower in HFD-fed mice receiving 10-HSA than in HFD controls (Supplementary Fig. 6d). Furthermore, 10-HSA supplementation significantly reduced serum total cholesterol and low-density lipoprotein cholesterol (LDL-C) levels, whereas the reduction in triglycerides was modest and did not reach statistical significance (Welch’s t-test; Fig. 6h, i). Although high-density lipoprotein cholesterol (HDL-C) levels were also decreased, the LDL-C/HDL-C ratio was 13.4% lower in 10-HSA-treated HFD-fed mice than in HFD-fed mice (Fig. 6i). These findings indicate that the metabolic effects of 10-HSA extend beyond glucose homeostasis to circulating lipid profiles and markers of HFD-induced hepatic injury.

Finally, we examined whether these metabolic phenotype improvements were accompanied by reduced hepatic lipid accumulation. Hematoxylin and eosin (H&E) and Oil Red O staining revealed pronounced hepatic pathological changes characterized by steatosis and substantial lipid deposition in HFD-fed mice. Quantitative analysis showed that 10-HSA supplementation reduced the hepatic histopathological score and Oil Red O-positive area by 30.8% and 81.0%, respectively (Fig. 6j–m). These histological findings were consistent with the improvements in circulating biochemical markers observed following 10-HSA treatment.

Together, these findings demonstrate that 10-HSA protects against multiple features of HFD-induced metabolic dysfunction, including excessive body-weight gain, impaired glucose homeostasis and insulin sensitivity, dyslipidemia, hepatic injury and steatosis. These *in vivo* findings demonstrate that 10-HSA is metabolically active and provide functional support for an OhyA**–**10-HSA axis downstream of *R. hominis*.

### 10-HSA restores hepatic lipid homeostasis through PPARα-associated transcriptional and metabolic reprogramming in HFD-fed mice

Given the metabolic benefits of 10-HSA, we first asked whether these effects were associated with remodeling of the gut microbiota; we first analyzed fecal metagenomic profiles across the four experimental groups (Methods). HFD feeding markedly altered gut microbial community structure, whereas 10-HSA supplementation produced comparatively modest changes and did not substantially reverse the HFD-associated microbial configuration (Supplementary Fig. 7a-c). These findings argue against broad restructuring of the gut microbiota as the primary mechanism underlying the metabolic effects of 10-HSA and prompted us to investigate its effects on host metabolic pathways.

We therefore profiled the hepatic transcriptional response to 10-HSA by RNA sequencing (Fig. 7a and Methods). PCA revealed a pronounced shift in liver transcriptomic profiles following HFD feeding, with HFD + 10-HSA mice separating from untreated HFD controls (Fig. 7b). Differential-expression analysis identified coordinated changes in genes involved in fatty-acid uptake, activation, desaturation and oxidation (Fig. 7c). Consistent with these changes, pathway analysis revealed enrichment of peroxisome proliferator-activated receptor (PPAR) signaling, fatty-acid degradation, linoleic acid metabolism, unsaturated fatty-acid biosynthesis and cytochrome P450-mediated lipid metabolism (Fig. 7d). Among the affected genes, *Ppara* and several established components of PPARα-regulated lipid metabolism genes, including *Cyp4a14*, *Hmgcs2*, *Acsl1*, and *Scd1*, were upregulated following 10-HSA supplementation. To determine whether selected transcriptional changes were reflected at the protein level, we examined PPARα and representative proteins involved in hepatic lipid metabolism (Methods). Western blotting showed increased hepatic abundance of PPARα, ACSL1, and HMGCS2 following 10-HSA supplementation (Fig. 7e,f). The increase in ACSL1 is consistent with enhanced activation of long-chain fatty acids for subsequent metabolic utilization, whereas elevated HMGCS2 suggests increased hepatic ketogenic potential. Notably, despite the transcriptional upregulation of *Scd1* and *Cyp4a14*, the abundance of their corresponding proteins was reduced (Supplementary Fig. 8a-c), indicating that these endoplasmic reticulum-associated enzymes may be subject to additional regulation at the translational or post-translational level. Taken together, these findings indicate that 10-HSA induces coordinated remodeling of hepatic lipid metabolism, characterized by activation of PPARα-associated pathways together with enhanced fatty-acid handling and ketogenic capacity.

**Figure 7:**
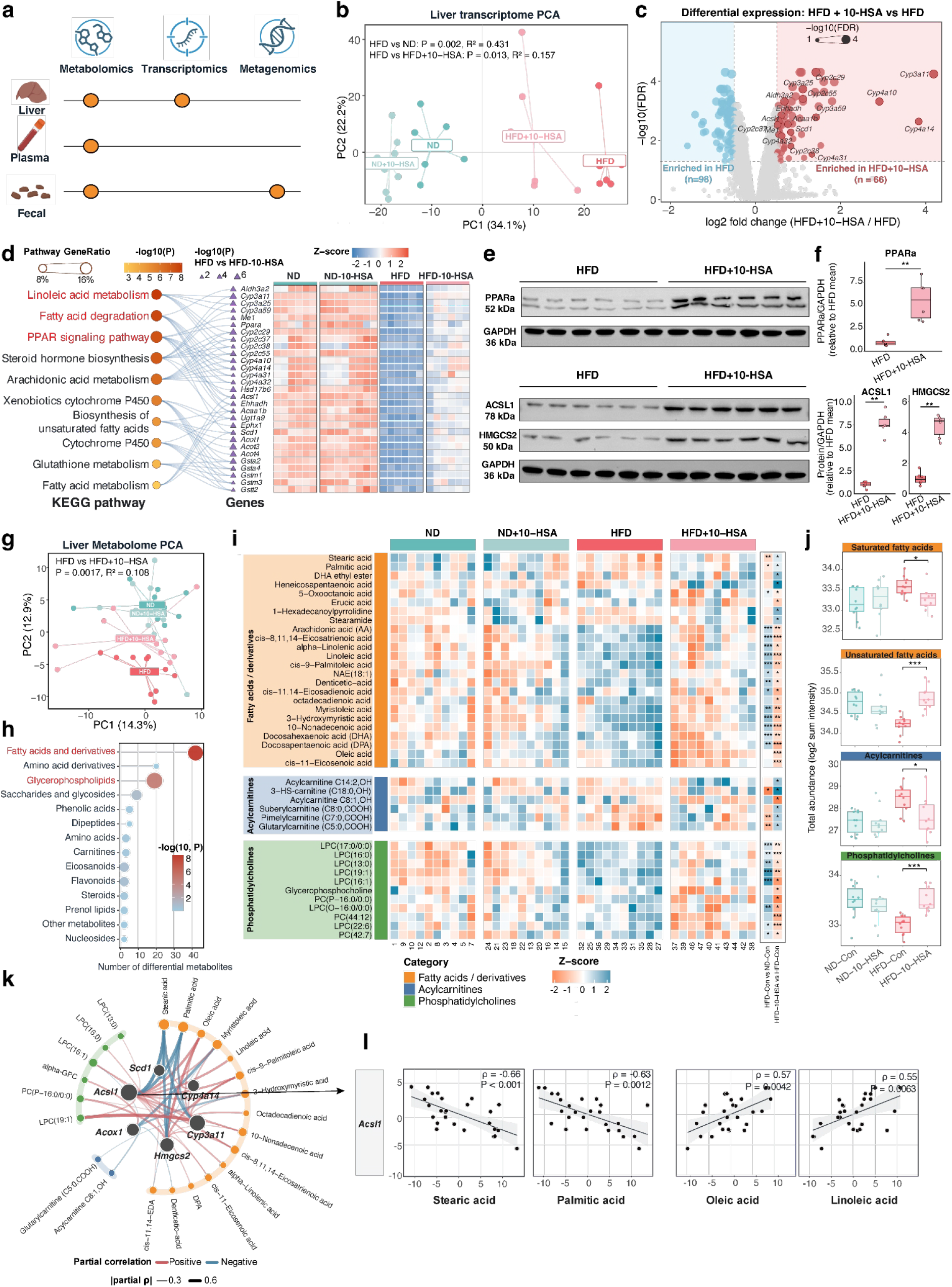
10-HSA alleviates HFD-induced hepatic lipid dysregulation through PPARα-associated metabolic reprogramming. (a) Overview of sample collection and multi-omics profiling, including metagenomic, transcriptomic, and metabolomic analyses. (b) PCA of liver transcriptomic profiles across the four experimental groups. (c) Differential gene expression between HFD and HFD + 10-HSA mice. (d) KEGG pathway enrichment of differentially expressed genes and expression patterns of genes contributing to the enriched pathways across the four experimental groups. (e) Representative western blots of hepatic PPARα, ACSL1, HMGCS2, and GAPDH (loading control). (f) Densitometric quantification of the indicated proteins normalized to the corresponding loading control. Each dot represents an individual mouse; bars indicate the mean ± s.e.m. *n* = 6 mice per group. (g) PCA was based on lipid-related metabolites (n = 10 mice per group). (h) Chemical-class distribution of differential hepatic metabolites between HFD and HFD + 10-HSA mice. (i) Heatmaps showing the relative abundance of representative hepatic fatty acids and derivatives, acylcarnitines, and phosphatidylcholines across the four experimental groups. (j) Aggregate abundance of saturated fatty acids, unsaturated fatty acids, acylcarnitines, and phosphatidylcholines across the four experimental groups. (k) Integrated correlation network linking PPARα-associated lipid-metabolism genes with differential hepatic metabolites. (l) Partial Spearman correlations between hepatic *Acsl1* expression and fatty acid abundance, adjusted for the four treatment groups; shaded areas indicate 95% confidence intervals, and P values are shown.

To determine whether this molecular program was accompanied by corresponding changes in lipid metabolism, we performed metabolomic profiling of liver, plasma, and fecal samples. Among the compartments examined, the liver exhibited the most pronounced metabolic response to 10-HSA, whereas changes in plasma and fecal metabolomes were comparatively modest (Supplementary Fig. 7d-f), highlighting the liver as a prominent site of metabolic response to 10-HSA. Consistently, hepatic metabolomic profiles clearly separated HFD + 10-HSA mice from untreated HFD controls (Fig. 7g). Differential metabolites were predominantly fatty acids and their derivatives, acylcarnitines, and glycerophospholipids (Fig. 7h), closely paralleling the lipid-metabolic pathways identified by transcriptomic analysis.

At the metabolite level, 10-HSA supplementation reduced hepatic levels of several saturated fatty acids, including palmitic and stearic acids, while increasing unsaturated fatty acids, including oleic and linoleic acids (Fig. 7i, j). This shift in hepatic fatty-acid composition was consistent with the observed changes in genes involved in fatty-acid desaturation and lipid remodeling. Hepatic acylcarnitines were also reduced following 10-HSA supplementation. Given that acylcarnitine accumulation can accompany an imbalance between fatty-acid delivery and oxidative capacity^32^, their reduction was consistent with improved hepatic fatty-acid handling. In parallel, several phosphatidylcholines and lysophosphatidylcholines increased following 10-HSA treatment, indicating substantial remodeling of hepatic glycerophospholipid metabolism. Together, these metabolomic changes were concordant with the PPARα-associated transcriptional and protein-level program induced by 10-HSA.

Integration of the hepatic transcriptomic and metabolomic datasets further revealed coordinated relationships between lipid-metabolism genes and altered metabolites (Fig. 7k and Methods). Notably, *Acsl1* expression was negatively correlated with hepatic stearic and palmitic acid levels but positively correlated with oleic and linoleic acid levels (Fig. 7l). Mechanistically, this correlation pattern is consistent with coordinated long-chain fatty-acid activation and desaturation. ACSL1 protein activates palmitate, stearate, oleate, and linoleate to their corresponding acyl-CoAs^33^ and channels these activated fatty acids toward β-oxidation or esterification into complex lipids^34,35^. Accordingly, the inverse associations of *Acsl1* expression with free palmitate and stearate may reflect enhanced acyl-CoA formation and downstream utilization of saturated fatty acids, consistent with evidence that hepatocyte-specific ACSL1 deficiency reduces fatty acyl-CoA availability and promotes hepatic free-fatty-acid accumulation^36^.

Taken together, the concordant transcriptomic, protein, and metabolomic responses support a liver-centered model in which 10-HSA induces PPARα-associated lipid-metabolic reprogramming. This program is characterized by coordinated changes in fatty-acid oxidation, activation, and desaturation, together with reduced accumulation of saturated fatty acids and acylcarnitines and remodeling of hepatic glycerophospholipids. These molecular and metabolic changes provide a mechanistic framework for the reduced hepatic steatosis and broader improvements in metabolic function observed in 10-HSA-treated HFD-fed mice.

## Discussion

Our study identifies a diet-responsive *R. hominis**–***OhyA***–***10-HSA axis linking gut microbial ecology and strain-level functional variation to host metabolic health. Across independent cohorts representing metabolic disorders and dietary intervention, *R. hominis* was consistently decreased in metabolic disorders and increased in dietary intervention. We then connected this human association to a defined microbial function by strain-level genetic variation, and metabolomic analyses further prioritized the oleate hydratase gene *ohyA* as a functional determinant of *R. hominis* metabolic activity. Wild-type OhyA protein converted oleic acid to 10-hydroxystearic acid (10-HSA), whereas an OhyA variant carrying five missense substitutions showed markedly impaired enzymatic activity. In HFD-fed mice, 10-HSA supplementation alleviated systemic metabolic phenotypes and restored hepatic lipid metabolism involving PPARα-associated fatty-acid metabolic pathways. Together, these findings provide a mechanistic and translational starting point for developing microbiome-informed interventions for metabolic disease.

Consistent with our findings, an independent study also reported that oral administration of *R. hominis* ameliorated obesity-related metabolic phenotypes in HFD-fed mice, further supporting a potentially beneficial role of this species in metabolic health^37^. Our multi-cohort human analyses complement these observations by showing that *R. hominis* abundance is reproducibly associated with metabolic health across both disease and dietary intervention settings. Importantly, we connect it to a specific lipid-metabolic enzyme and to a host-active metabolite, suggesting that differences in the functional capacity of *R. hominis* may contribute to variation in its metabolic effects across individuals. Thus, this shifts the interpretation of *R. hominis* from a descriptive microbial marker to a tractable entry point for metabolic intervention. And the therapeutic strategies could target the bacterium itself, its enzyme-mediated metabolic capacity, or the metabolites generated from dietary substrates.

The ecological role of OhyA may nevertheless originate primarily from microbial adaptation rather than host benefit. Although OhyA proteins are distributed across phylogenetically diverse bacteria and differ in catalytic efficiency, substrate preference, and product specificity^38,39^, the structural and mechanistic basis of OhyA function has been studied primarily using enzymes from *S. aureus* and *E. meningoseptica*^40,41^, whereas *R. hominis* OhyA remains poorly characterized. In several bacteria, OhyA-mediated hydration reduces the antimicrobial activity of unsaturated fatty acids and may therefore support survival in fatty-acid-rich environments^42,43^. The effects of 10-HSA on host metabolism may thus represent a physiologically relevant cross-kingdom consequence of a bacterial stress-adaptation pathway. Our finding that five amino-acid substitutions impair the catalytic activity of *R. hominis* OhyA provides an entry point for developing OhyA variants and *R. hominis* strains with improved catalytic efficiency, stability, or substrate breadth.

The biological relevance of 10-HSA can be considered within a broader network of microbial fatty-acid metabolites. Oleic acid and linoleic acid are abundant dietary C18 unsaturated fatty acids that can be transformed by gut bacteria into a series of hydroxy, oxo, conjugated, and partially saturated fatty-acid intermediates, several of which can exert distinct effects on the host^44^. For example, the linoleic-acid-derived microbial metabolite 10-hydroxy-cis-12-octadecenoic acid improves intestinal barrier integrity^45^, while 10-oxostearic acid, another microbial oleic-acid-derived metabolite, directly binds and activates PPARα and protects mice against colitis^46^. Work on *S. aureus* OhyA-derived hydroxy fatty acids was reported to activate PPARα in macrophages during skin infection, linking this bacterial pathway to host immune modulation^47^. Separately, 10-HSA has also been shown to activate PPARα in intestinal epithelial cells and improve hepatic lipid metabolism in a liver-injury mouse model, respectively^48,49^. In our HFD model, 10-HSA treatment was accompanied by coordinated remodeling of hepatic fatty-acid metabolism, including transcriptional enrichment of PPARα-associated pathways and increased hepatic abundance of ACSL1 and HMGCS2. The discordant regulation of *Scd1* and *Cyp4a14* at the transcript and protein levels suggests that their protein abundance may be shaped by regulatory processes beyond transcription. This is particularly plausible for SCD1, a short-lived endoplasmic-reticulum membrane protein known to undergo rapid post-translational turnover, which can uncouple changes in *Scd1* mRNA expression from steady-state protein abundance^50,51^. These observations provide a basis for investigating microbial metabolite-host interaction, particularly the 10-HSA–PPARα-associated hepatic lipid-metabolic response, in metabolic disorders.

Diet may provide an important ecological context for this interaction. Oleic and linoleic acids are not only absorbed and metabolized by the host but also represent substrates and potential selective pressures for intestinal bacteria capable of transforming unsaturated fatty acids. Mediterranean-style dietary patterns, which commonly contain foods rich in unsaturated fats, are associated with distinct microbial taxonomic and functional configurations, and their cardiometabolic effects can vary according to baseline microbiome composition^52^. Diet has also been associated with the distribution of functionally distinct strains within individual bacterial species^53^. Our observation that *R. hominis* abundance increased in distinct dietary interventions, together with longitudinal shifts in the *ohyA* allelic profile, raises the possibility that dietary environments influence both the abundance and functional composition of *R. hominis*. Such changes should not be interpreted as rapid reversal or acquisition of individual SNPs; rather, they may reflect changes in the relative representation of pre-existing strains carrying different *ohyA* alleles. These raise the possibility that dietary lipids could be used to support or complement microbial pathways that generate host-active metabolites, whereas individuals with low *R. hominis* abundance or OhyA activity may require direct metabolite supplementation or defined microbial interventions.

Several limitations should be considered. First, although the associations among *R. hominis*, microbial genetic variation, circulating metabolites, and cardiometabolic traits were supported by multiple human cohorts and complementary analyses, much of the human evidence remains observational. The SNP-metabolite-trait mediation analyses should therefore be interpreted as statistical support for potential intermediate metabolic pathways rather than evidence of biological causality. Second, the detailed strain-level genetic associations and SNP-defined participant stratification were primarily established in the MetaCardis cohort. Although the longitudinal FBIP cohort provided complementary evidence that the *ohyA* allelic profile changes during dietary intervention, independent replication of the genetic signatures in additional deeply phenotyped cohorts will be important. Moreover, longitudinal changes in metagenomic *ohyA* profiles probably reflect shifts in the relative representation of *R. hominis* strains rather than de novo mutation or reversal of individual variants; strain-resolved longitudinal sequencing and isolate-based studies will be needed to distinguish strain replacement from within-strain evolution. Third, although the combined five-variant OhyA protein showed markedly impaired enzymatic activity, our experiments do not resolve the individual contribution of each amino-acid substitution or potential interactions among them. Finally, our liver multi-omics data strongly implicate PPARα-associated lipid metabolism in the response to 10-HSA but do not establish that PPARα is required for its metabolic effects.

Future studies integrating strain-resolved longitudinal metagenomics, species-resolved metatranscriptomics, and stable-isotope tracing with labeled oleic acid could further define which intestinal strains actively produce 10-HSA and trace microbial carbon from the gut into the circulation and host tissues. PPARα perturbation, direct ligand-binding and transcriptional activation assays will clarify the host receptor mechanism, and profiling of additional OhyA products will determine whether 10-HSA is the principal effector or part of a broader metabolite network. Ultimately, these studies could support a progression from biomarker discovery to defined strains, enzymes, or metabolites designed to improve metabolic health.

Together, our findings support a therapeutic framework in which the gut microbiome is targeted not only by altering community composition, but also by restoring or enhancing microbial functions and their host-active metabolites. The diet-responsive *R. hominis*–OhyA pathway provides a concrete example of this approach, in which a dietary fatty acid is converted by a functionally variable microbial enzyme into 10-HSA, which improves metabolic phenotypes *in vivo.* This work provides a foundation for evaluating high-activity *R. hominis* strains, engineered OhyA variants, and 10-HSA or related metabolites as complementary strategies for metabolic disorders.

## Methods

### Study cohorts

We integrated five independent human cohorts comprising 924 participants to investigate gut microbial features associated with metabolic disorders and responses to dietary interventions designed to improve metabolic health (Fig. 1a and Supplementary Table 1).

Two cross-sectional cohorts were included. The Suzhou cohort comprised 253 participants from Suzhou, China, including 96 individuals with normal glucose tolerance (NGT), 80 individuals with prediabetes, and 77 individuals with type 2 diabetes (T2D). The MetaCardis cohort comprised 429 participants from a European multicenter study, including 270 healthy controls and 159 untreated metabolically matched controls (UMMC), individuals with features of the metabolic syndrome who were not receiving lipid-lowering, antidiabetic, or antihypertensive medication.

Three longitudinal dietary intervention cohorts included individuals with prediabetes. The fiber-based intervention program (FBIP), conducted in Singapore, included 54 participants who underwent a 16-week legume-enriched dietary intervention. Fecal samples were collected at baseline and at weeks 2, 4, 8, 12, and 16. The personalized postprandial glucose-targeting diet (PPT) cohort included 92 participants who received a six-month personalized dietary intervention, whereas the Mediterranean diet (MED) cohort included 96 participants who followed a six-month Mediterranean dietary intervention. Both the PPT and MED cohorts were conducted in Israel, with samples collected at baseline and at the end of the intervention.

### Metagenomic processing and taxonomic profiling

Raw shotgun metagenomic reads were processed using fastp^54^ (v1.0.1) to remove adapter sequences, low-quality bases, and reads containing excessive ambiguous nucleotides. Quality-filtered reads were aligned to the human reference sequence hg37dec_v0.1 using Bowtie2^55^ (v2.5.4), and reads mapping to the human genome were removed. The remaining non-host reads were retained for microbial taxonomic profiling.

Taxonomic profiles were generated using MetaPhlAn^56^ (v4.2.2) with the CHOCOPhlAn species-level genome bin marker database (mpa_vJan25_CHOCOPhlAnSGB_202503). Duplicate taxonomic entries were removed before downstream analysis. Taxa were retained if detected in at least 10% of samples and with a relative abundance of at least 0.0002 when present; taxa not meeting these criteria were excluded.

### Differential abundance analysis and candidate-taxon prioritization

Statistical analyses were performed in R (v4.5.1) unless otherwise specified. Between-group differences in the two cross-sectional cohorts were evaluated using the Wilcoxon rank-sum test. In the Suzhou cohort, participants with prediabetes were compared with NGT participants, whereas the T2D group was included for descriptive visualization. In the MetaCardis cohort, healthy controls were compared with UMMC participants. Taxa with *P* < 0.05 and lower relative abundance in the metabolically disordered group in both cohorts were retained as metabolic disorder-associated candidates.

Within-participant changes in the three dietary intervention cohorts were evaluated using the paired Wilcoxon signed-rank test. In the FBIP cohort, candidate selection was based on the comparison between baseline and week 16; comparisons between baseline and weeks 2, 4, 8, and 12 were used to characterize temporal changes. In the PPT and MED cohorts, baseline samples were compared with samples collected at the end of the intervention. Taxa with *P* < 0.05 and increased relative abundance in all three intervention cohorts were retained as diet-responsive candidates. The metabolic disorder-associated and diet-responsive candidate sets were intersected to prioritize taxa for downstream strain-level and functional analyses.

### Microbial single-nucleotide polymorphism (SNP) identification and genotype processing

To characterize within-species genetic variation in *R. hominis*, quality-filtered non-host reads were aligned to the reference genome GCA_902387955.1 (Unified Human Gastrointestinal Genome identifier: MGYG-HGUT-02517) using Bowtie2. Alignments were sorted and filtered using SAMtools^57^ (v1.22), retaining primary alignments with mapping quality scores of at least 20. Single-nucleotide polymorphisms (SNPs) were jointly identified across samples using SAMtools mpileup followed by BCFtools call^58^ (v1.22), with minimum mapping and base quality thresholds of 20 and 25, respectively. Only biallelic SNPs were retained for downstream analyses. At each retained locus, samples were classified as carrying either the reference genotype (0) or the non-reference genotype (1) according to the genotype with the lower Phred-scaled likelihood value, PL(0/0) or PL(1/1).

### Microbial genome-wide association (GWAS) and gene-level enrichment analyses

Microbial genome-wide association analyses were performed using PLINK^59^ (v1.9) based on the processed *R. hominis* SNP genotype matrix. Linear regression models were used to test associations between individual SNPs and eight host cardiometabolic traits: glycated hemoglobin (HbA1c), body mass index (BMI), waist circumference, fasting plasma triglycerides, fasting plasma adiponectin, fasting plasma C-reactive protein (CRP), systolic blood pressure, and diastolic blood pressure. Country, sex, age, and the relative abundance of *R. hominis* were included as covariates. SNPs with a minor allele frequency below 0.01 were excluded from the analyses.

All filtered SNPs were mapped to annotated genes in the *R. hominis* reference genome. For each cardiometabolic trait, SNPs were ranked according to their association *P* values, and those within the top 1% were defined as top-ranked association signals. SNPs ranked within the top 1% for at least seven of the eight phenotypes were mapped to genes and used for Gene Ontology enrichment analysis^26^. Separately, for each gene, a hypergeometric test was used to determine whether top-ranked SNPs were overrepresented relative to the total number of filtered SNPs mapped to that gene, using all filtered SNPs as the background. To assess the robustness of the gene-level enrichment, we additionally performed 10,000 SNP-level permutations by randomly reassigning the observed top-ranked SNP profiles across the 97,252 eligible SNP positions while preserving the phenotype-specific SNP counts and cross-phenotype overlap structure. Gene-level enrichment was recalculated for each permutation to generate empirical *P* values, which were adjusted using the Benjamini-Hochberg procedure. Genes showing significant enrichment (FDR-adjusted *P* < 0.05) and containing top-ranked association signals across all eight cardiometabolic traits were retained as candidate genes for downstream analyses.

### SNP-based classification and genetic structure analysis

To assess whether *R. hominis* genetic variation provided information beyond species-level abundance, random forest models were constructed using the 603 SNPs located within the 68 genes showing recurrent top-ranked association signals across seven or eight cardiometabolic traits. Four models were evaluated: host covariates alone (country, sex, and age), host covariates plus *R. hominis* relative abundance, host covariates plus SNP profiles, and host covariates plus both relative abundance and SNP profiles. Random forests were fitted with 1,000 trees using class-balanced sampling, and model discrimination was evaluated from out-of-bag predictions using receiver operating characteristic curves and the area under the curve (AUC). To further characterize differences in overall SNP profiles between healthy controls and UMMC participants, SNPs with more than 50% missing genotypes were excluded, and participants with genotype call rates below 50% were removed. Missing genotypes were median-imputed, and principal component analysis (PCA) was performed on the centered and scaled binary genotype matrix after removing invariant SNPs. Group differences in SNP profiles were assessed by PERMANOVA based on Euclidean distances with 999 permutations.

### SNP-based phylogenetic reconstruction and UMMC enrichment analysis

The 85 SNPs located within the six candidate genes were converted into a nucleotide alignment using VCF2PHYLIP^60^ (v2.9). A maximum-likelihood tree was reconstructed with IQ-TREE^61^ (v2.4.0) under the GTR+G model, and branch support was assessed using 1,000 ultrafast bootstrap replicates and 1,000 SH-like approximate likelihood-ratio test replicates. Samples were ordered according to the circular tree topology. A sliding-window analysis with a 30° window and 5° step was then used to identify UMMC-enriched segments. For each window, UMMC enrichment relative to the overall cohort proportion was tested using a one-sided hypergeometric test with Bonferroni correction. Overlapping significant windows were merged into continuous enriched segments, with the remaining tree positions defined as background. Among UMMC participants, non-reference genotype frequencies across the 85 candidate SNPs were compared between enriched and background segments using Fisher’s exact tests with 10,000 Monte Carlo simulations and Benjamini-Hochberg correction. The eight cardiometabolic traits were then compared between the same two groups using Wilcoxon rank-sum tests.

### SNP-metabolite association and mediation analyses

The 85 candidate SNPs were tested for associations with 1,513 serum metabolites in participants with matched genotype and metabolomics data. Metabolite origins were classified using the metabolite source database implemented in TidyMass2^62,63^. Serum metabolite levels were compared between reference and non-reference genotype groups using Welch’s two-sample *t*-test. Effect sizes were summarized as mean differences and linear-model R^2^ values, and association *P* values were adjusted using Bonferroni correction. The six candidate genes were ranked according to the number of significant SNP-metabolite associations involving SNPs within each gene.

Mediation analyses were performed for selected SNP-metabolite-phenotype relationships using the *mediation* package^64^ (v4.5.1) in R. Missing values in the selected metabolite and phenotype variables were imputed using the corresponding median. SNP genotype, serum metabolite, and cardiometabolic traits were specified as the exposure, mediator, and outcome, respectively. Linear mediator and outcome models were fitted without additional covariates. Average causal mediation effects and 95% confidence intervals were estimated using 500 nonparametric bootstrap simulations, and the resulting *P* values were adjusted using the Benjamini-Hochberg method.

### Construction of the *ohyA*-derived genetic burden score

To summarize the cumulative contribution of genetic variation within *ohyA*, we constructed a derived genetic burden score (dGBS) using the retained SNPs within this gene. A metabolic syndrome-related cardiometabolic trait axis was constructed from four equally weighted binary components: central obesity, defined as a waist circumference of at least 94 cm in men or 80 cm in women; elevated triglycerides, defined as at least 1.7 mmol/L; elevated blood pressure, defined as systolic blood pressure of at least 130 mmHg or diastolic blood pressure of at least 85 mmHg; and elevated glycaemia, defined as HbA1c of at least 5.7%. The four components were summed to produce a score ranging from 0 to 4 and subsequently standardized as a z-score.

In the MetaCardis cohort, two complementary dGBS measures were generated. The group-derived dGBS was weighted by SNP effect estimates from logistic regression comparing UMMC participants with healthy controls, whereas the phenotype-derived dGBS was weighted by effect estimates from linear regression using the standardized cardiometabolic trait axis as the outcome. Both models included age and sex as covariates.

For each individual, the dGBS was calculated as:

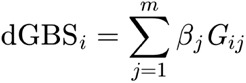

where Gᵢ_ⱼ_ denotes the genotype of SNP j in individual i, coded as 0 for the reference genotype and 1 for the non-reference genotype; β_ⱼ_ denotes the corresponding regression coefficient; and m denotes the number of SNPs included in the score. The resulting dGBS values were standardized as z-scores. Associations between the phenotype-derived dGBS and individual cardiometabolic traits were subsequently evaluated. The same phenotype-axis framework was independently applied to baseline samples from the FBIP cohort to derive a cohort-specific phenotype-derived dGBS. Group differences in dGBS were evaluated using the Wilcoxon rank-sum test. Correlations between the group-derived and phenotype-derived dGBS measures were assessed using Pearson correlation, whereas associations between the phenotype-derived dGBS and individual cardiometabolic traits were assessed using Spearman correlation. Age- and sex-adjusted linear models were used to evaluate dGBS-by-group interactions.

### Identification and functional prediction of *ohyA* missense variants

Variants within the *ohyA* coding region were annotated as synonymous or missense substitutions based on the reference genome sequence and gene annotation. Missense variants were evaluated using the DDMut web server (accessed October 2025)^30^ and PROVEAN (v1.1.5)^31^ to predict their effects on protein stability and function, respectively. Variants predicted to destabilize OhyA by DDMut and to be deleterious by PROVEAN were prioritized as candidate functional variants and mapped onto the predicted OhyA protein structure.

### *R. hominis* culture and oleic-acid incubation assay

*R. hominis* DSM16839 was routinely cultured in a vinyl anaerobic chamber (75% N_2_/20% CO_2_/5% H_2_). 50 μL of overnight culture was added in triplicate to Hungate tubes containing 5 mL of basal medium (medium composition details in Supplementary Table 4), basal medium supplemented with ethanol (vehicle control), and basal medium supplemented with 1 mM oleic acid. The tubes were then incubated at 37 °C overnight, followed by centrifugation at 3,000 × g for 10 minutes and stored at - 80 °C for further analysis.

### LC-MS detection and quantification of OA and 10-HSA

This experiment was conducted at the Singapore Phenome Center (SPC), Lee Kong Chian School of Medicine, Nanyang Technological University, Singapore. For targeted LC-MS analysis of the *in vitro* samples, aliquots of the collected supernatants were extracted with methanol (CAS 67-56-1; Merck, Germany) containing 0.1% formic acid (CAS 64-18-6; Fisher Chemical, USA). Samples were sonicated in an ice-water bath for 10 min, incubated on ice for an additional 10 min, and centrifuged at 14,000 × *g* for 15 min at 4 °C. The resulting supernatants were collected and stored at -80 °C until analysis.

A 10 μL aliquot of each sample was subjected to targeted LC-MS analysis using an AB Sciex QTRAP 6500 mass spectrometer (SCIEX). OA (Sigma, O1008) and 10-HSA (MCE, HY-W128156) were identified and quantified using authentic reference standards and calibration curves. Data acquisition, peak integration, metabolite quantification, and quality assessment were performed using Analyst software (SCIEX, USA). Quality-control samples were included throughout the analytical batch to monitor analytical reproducibility and instrument performance. Metabolite concentrations were reported in μM.

### Recombinant expression and purification of wild-type and 5-substitution OhyA

The coding sequence of the *R. hominis ohyA* gene was retrieved from the NCBI database and commercially synthesized by Axil Scientific Pte. Ltd. (Singapore). The wild-type sequence and a variant carrying the five prioritized missense substitutions were codon-optimized for expression in *Escherichia coli* and engineered with an N-terminal 6×His tag. The resulting *ohyA*-WT and *ohyA*-5-SNP constructs were cloned into the pET-28a (+) expression vector for recombinant protein production.

Protein expression was induced with 0.5 mM isopropyl β-D-1-thiogalactopyranoside (IPTG) at 20 °C with shaking overnight, and all the following protein purification procedures were performed on ice or at 4°C. Cells were harvested by centrifugation and lysed by sonication in a PBS buffer (pH 7.2, Vivantis, PB0342) containing 1 mM phenylmethylsulfonyl fluoride (PMSF). The cell lysates were centrifuged at 33,000 × g for 30 min, and the resulting supernatants were subjected to Ni²⁺-affinity chromatography. The nickel column (Cytiva Life Sciences, 17524801) was washed with 2-5 column volumes of PBS containing 25 mM imidazole to remove nonspecifically bound proteins, followed by elution of the recombinant proteins with PBS containing 100 mM imidazole. The eluted proteins were further purified by size-exclusion chromatography using a Superdex-200 column (Cytiva Life Sciences, 28989335) equilibrated with PBS. Protein-containing fractions were collected, and the expression and purity of recombinant wild-type OhyA (OhyA-WT) and the five-substitution mutant (OhyA-5-SNP) were assessed by SDS-PAGE and stained with Coomassie blue dye. The accurate molecular masses of the purified proteins were determined by matrix-assisted laser desorption/ionization time-of-flight mass spectrometry (MALDI-TOF MS), and their identities were further confirmed by de novo sequencing analysis.

### Cell-free OhyA enzymatic assay

The substrate-conversion activities of recombinant OhyA-WT and OhyA-5-SNP were assessed by liquid chromatography–mass spectrometry (LC-MS) by measuring the conversion of oleic acid (OA) to 10-hydroxystearic acid (10-HSA). Each enzymatic reaction was performed in a total volume of 200 μL containing 0.25 mg/mL purified OhyA-WT or OhyA-5-SNP, 50 mM potassium phosphate buffer (pH 6.0), 10 mM sodium chloride (NaCl; S9888), 10 mM dithiothreitol (DTT; D9163), 50 μM flavin adenine dinucleotide (FAD; F6625), 0.2 mg/mL bovine serum albumin (BSA; B4287), and 1 mg/mL OA (O1008). All reagents listed above were purchased from Sigma-Aldrich (St. Louis, MO, USA). Reaction mixtures were incubated at 37 °C for 24 h, after which the reactions were terminated by the addition of 80 μL acetonitrile. Samples were centrifuged at 4,000 × *g* for 10 min, and the resulting supernatants containing OA and the 10-HSA reaction product were collected for LC-MS analysis.

### Mouse model and 10-HSA administration

Male specific pathogen-free (SPF) C57BL/6J mice aged 6 weeks were obtained from Shanghai SLAC Laboratory Animal Co., Ltd. (Shanghai, China). After one week of acclimatization, mice were randomly assigned to four experimental groups (n = 10 per group): normal diet control (ND), normal diet with 10-HSA treatment (ND+10-HSA), HFD, and high-fat diet with 10-HSA treatment (HFD+10-HSA). Mice in the ND groups were fed a standard laboratory chow diet (catalog no. 101001, Xietong Bioengineering, China), whereas mice in the HFD groups were fed a high-fat diet providing 60% of total energy from fat (D12492, Research Diets, Inc., New Brunswick, NJ, USA). Mice in the ND + 10-HSA and HFD + 10-HSA groups were administered 10-HSA by oral gavage at 100 mg/kg body weight/day throughout the 8-week intervention^49^. Body weight was monitored throughout the study. At the experimental endpoint, mice were fasted for 6-8 h before sample collection. All animal procedures were conducted in accordance with institutional guidelines and were approved by the Experimental Animal Welfare and Ethics Committee of Nanjing Medical University (IACUC no. IACUC-260507).

### Glucose and insulin tolerance tests

Glucose tolerance tests (GTTs) and insulin tolerance tests (ITTs) were performed during week 7 of the dietary intervention. The GTT was conducted first, followed by the ITT 2 days later. For the GTT, food was removed overnight before the test, and mice remained fasted throughout the experiment. Baseline blood glucose was measured from tail blood immediately before glucose administration (0 min). Mice were then administered glucose (1.5 g/kg body weight) by intraperitoneal injection. Blood glucose concentrations were measured at 15, 30, 60, and 120 min after glucose administration. For the ITT, mice were fasted for 4 h before the test and remained fasted throughout the experiment. Baseline blood glucose was measured from tail blood immediately before insulin administration (0 min). Mice were then administered insulin (1.0 IU/kg body weight) by intraperitoneal injection, and blood glucose concentrations were measured at 15, 30, 60, and 120 min after injection. Glucose tolerance and insulin sensitivity were evaluated from the resulting blood-glucose response curves, and the AUC was calculated for each mouse.

### Fasting metabolic measurements and serum biochemistry

At the experimental endpoint, mice were fasted for 6-8 h before blood collection. Whole blood was collected and allowed to clot, followed by centrifugation to obtain serum. Serum biochemical parameters, including alanine aminotransferase (ALT), aspartate aminotransferase (AST), total cholesterol (TC), triglycerides (TG), high-density lipoprotein cholesterol (HDL-C), and low-density lipoprotein cholesterol (LDL-C), were measured using an automated clinical chemistry analyzer according to the manufacturer’s instructions. Fasting blood glucose was measured using a glucometer. Fasting serum insulin concentrations were determined separately, and insulin resistance was estimated using the HOMA-IR.

### Liver and colon histology and hepatic lipid assessment

For hematoxylin and eosin (H&E) staining, liver and colon tissues were fixed in 4% paraformaldehyde for 48 h. Fixed tissues were dehydrated through a graded ethanol series (50%, 70%, 85%, 95%, and 100%), cleared twice in xylene for 5 min each, embedded in paraffin, and sectioned at a thickness of 4 μm. Paraffin sections were deparaffinized twice in xylene for 10 min each and rehydrated sequentially in 100%, 95%, 85%, and 75% ethanol for 5 min each, followed by rinsing in distilled water. Sections were stained with 2% hematoxylin for 5 min, differentiated in 1% hydrochloric acid in ethanol for 5 s, rinsed in running water for 1 min to allow bluing, and washed in distilled water for 1 min. Sections were subsequently counterstained with 0.5% eosin for 2 min, briefly rinsed in distilled water, dehydrated sequentially in 80% ethanol for 30 s, 95% ethanol for 2 min, and absolute ethanol for 3 min, and cleared in xylene for 3 min. Sections were mounted with neutral resin and examined using a light microscope. Liver sections were evaluated for hepatic steatosis and inflammatory changes, whereas colon sections were examined for overall mucosal and crypt architecture.

Hepatic lipid accumulation was assessed by Oil Red O staining of frozen liver sections. Fresh liver tissues were prepared for cryosectioning, and the resulting sections were air-dried before fixation in 4% paraformaldehyde for 15 min. Sections were washed in running water for 5 min and briefly equilibrated in 60% isopropanol before staining with Oil Red O working solution for 15 min. After staining, sections were briefly differentiated in 60% isopropanol and washed in running water for 2 min. Sections were counterstained with hematoxylin for 2 min, blued for 10 min, and mounted using glycerol-based mounting medium. Stained sections were examined and photographed under a light microscope. Hepatic lipid accumulation was quantified as the percentage of tissue area occupied by Oil Red O-positive lipid droplets.

### Mouse fecal metagenomic sequencing and microbiome analysis

Fresh fecal samples were immediately snap-frozen in liquid nitrogen and stored at -80 °C until processing. Metagenomic sequencing and primary bioinformatic processing were performed by Beijing Biomarker Technologies Co., Ltd. Total DNA was extracted using the TGuide S96 Magnetic Universal DNA Kit and the TGuide S96 Automated Nucleic Acid Extraction and Purification System (TIANGEN Biotech, Beijing, China). Briefly, samples were mixed with Buffers SA and SC and homogenized with eight 3-mm grinding beads and 0.2 g of 1-mm grinding beads using a TGrinder H24 Tissue Homogenizer (TIANGEN; OSE-TH-01) at 6 m/s for 20 s, followed by a 10 s interval, for two cycles. Samples were subsequently lysed at 70 °C, centrifuged, and the supernatants were processed with Buffer SH before automated DNA extraction and purification.

DNA concentration and purity were assessed using a NanoDrop 2000 spectrophotometer (Thermo Fisher Scientific) and a Qubit 3 fluorometer (Invitrogen), and DNA integrity was evaluated by agarose gel electrophoresis. Metagenomic libraries were constructed from 10 ng of qualified DNA using the VAHTS Universal Plus DNA Library Prep Kit for Illumina (Vazyme, ND617-02). DNA was enzymatically fragmented, end-repaired, and 3′-adenylated, followed by adaptor ligation, magnetic-bead purification, size selection and eight cycles of PCR amplification. Library products were purified using VAHTS DNA Clean Beads (Vazyme, N411-03). Library fragment size and concentration were assessed using a Qsep-400 system and Qubit 3 fluorometer, respectively. Qualified libraries, with concentrations ≥1 ng/μL and fragment-size center values of 430-530 bp, were sequenced on an Illumina NovaSeq X platform using a paired-end 150-bp strategy.

Raw reads were quality-filtered using fastp, and host-derived reads were removed by alignment against the Mus musculus reference genome (GRCm38, release 95) using Bowtie2. The resulting non-host clean reads were taxonomically classified using Kraken2^65^, and species-level abundances were estimated from read counts using Bracken^66^. Only taxa assigned to confirmed bacterial lineages were retained, whereas non-bacterial and unclassified taxa were excluded. Taxa detected in fewer than 12 of the 120 samples (< 10% prevalence) were removed, and the 1,000 most abundant bacterial species, ranked by total read abundance across all samples, were retained^67^. Shannon diversity was calculated from the rarefied count matrix using the *vegan* package^68^. Bray-Curtis dissimilarities were calculated from the same rarefied species-level matrix, and principal coordinate analysis was performed with Lingoes correction. Differences in microbial community composition among dietary groups at each time point were assessed by PERMANOVA with 9,999 permutations.

### Liver RNA sequencing and transcriptomic analysis

Total RNA was extracted from mouse liver tissues using TRIzol reagent (Life Technologies, Carlsbad, CA, USA). RNA concentration and purity were assessed using a NanoDrop 2000 spectrophotometer (Thermo Fisher Scientific), and RNA integrity was evaluated using the RNA Nano 6000 Assay Kit on an Agilent 2100 Bioanalyzer (Agilent Technologies). For each sample, 1 μg of total RNA was used for library preparation with the Hieff NGS Ultima Dual-mode mRNA Library Prep Kit for Illumina (Yeasen Biotechnology, Shanghai, China). Poly(A)-containing mRNA was enriched using oligo (dT)-conjugated magnetic beads, followed by cDNA synthesis, end repair, adaptor ligation, size selection, PCR amplification, and library quality assessment.

Libraries were sequenced on an Illumina NovaSeq platform to generate 150-bp paired-end reads. Raw reads were processed to remove adaptor-containing, poly-N, and low-quality reads, and sequencing quality was evaluated based on Q20, Q30, GC content, and sequence duplication rates. Clean reads were aligned to the mouse reference genome using HISAT2^69^, and transcript assembly and annotation were performed using StringTie. Gene expression was quantified as fragments per kilobase of transcript per million mapped fragments (FPKM).

PCA was performed with gene-wise centering without variance scaling. Differences between selected treatment groups were assessed by permutational multivariate analysis of variance based on Euclidean distances with 9,999 permutations, and unadjusted permutation-derived *P* values were reported. Differential expression between the HFD + 10-HSA and HFD groups was evaluated using six biological replicates per group. Genes with FPKM ≥ 1 in at least three of the 12 comparison samples were retained, and differential expression was analyzed on log2(FPKM + 1) values using the limma-trend framework with robust empirical Bayes moderation. Positive log2 fold-change values indicated higher expression in the HFD + 10-HSA group. *P* values were adjusted using the Benjamini-Hochberg procedure, and genes with a false discovery rate (FDR) < 0.05 and an absolute log2 fold change ≥ 0.5 were defined as differentially expressed. Gene Ontology and Kyoto Encyclopedia of Genes and Genomes pathway over-representation analyses were conducted using *clusterProfiler*^70^, with all genes tested in the differential-expression analysis as the background universe. Enrichment *P* values were adjusted using the Benjamini-Hochberg method, and terms with FDR < 0.05 were considered significantly enriched.

### LC-MS untargeted metabolomics of mouse liver, plasma and feces

LC-MS untargeted metabolomic profiling of mouse liver, plasma and feces was performed by Biotree Biomedical Technology (Shanghai, China) using a Vanquish UHPLC system coupled to an Orbitrap Exploris 120 high-resolution mass spectrometer (Thermo Fisher Scientific). Approximately 25 mg of liver or fecal material was extracted with 500 µL of pre-cooled (-40 °C) methanol/acetonitrile/water (2:2:1, v/v/v) containing the ISAMIDE and ISC18 internal-standard mixtures, homogenized at 35 Hz for 4 min and sonicated in an ice-water bath for 5 min for three cycles, incubated at -40 °C for 1 h, and filtered through a 96-well filtration plate; plasma (50 µL) was precipitated with 200 µL of pre-cooled methanol/acetonitrile (1:1, v/v) containing the same internal standards, shaken at 750 rpm for 5 min, allowed to stand for 5 min, and filtered. Pooled quality-control samples were prepared by combining equal aliquots of sample supernatants and were included to monitor analytical stability.

Polar metabolites were separated on an ACQUITY UPLC BEH Amide column (2.1 × 50 mm, 1.7 µm) using water containing 25 mmol/L ammonium acetate and 25 mmol/L ammonium hydroxide as mobile phase A and acetonitrile as mobile phase B, whereas non-polar metabolites were separated on a Kinetex C18 column (2.1 × 50 mm, 2.6 µm) using 0.01% acetic acid in water as mobile phase A and isopropanol/acetonitrile (1:1, v/v) as mobile phase B; the autosampler temperature was 4 °C and the injection volume was 2 µL. Data were acquired in positive- and negative-ion electrospray modes with information-dependent MS/MS acquisition using the following settings: sheath gas, 50 arbitrary units; auxiliary gas, 15 arbitrary units; capillary temperature, 320 °C; spray voltage, +3.8 kV or -3.4 kV; full-MS resolution, 60,000; MS/MS resolution, 15,000; and stepped normalized collision energies of 20, 30, and 40. Raw files were converted to mzXML using ProteoWizard^71^ (v3.0.24054), and feature detection, retention-time alignment, peak integration, and annotation were performed using an in-house R workflow based on XCMS^72^ and BiotreeDB v4.0.

Downstream analyses used level 1 metabolites, defined by concordant MS1 (accurate mass), MS/MS, and retention-time matching to authentic standards, from the provider-supplied peak-area matrix. For hepatic differential-metabolite analysis, HFD + 10-HSA and HFD mice were compared using two-sided Welch’s t-tests on log2-transformed intensities; effect sizes were expressed as differences in mean log2 intensity, and both nominal *P* values and Benjamini-Hochberg false-discovery rates were retained. Metabolites with nominal *P* < 0.05 were used for the chemical-class summary and lipid heatmap; class over-representation was assessed by a hypergeometric test against all retained level 1 liver metabolites, followed by Benjamini-Hochberg correction. For group-level lipid summaries, peak areas of the selected differential metabolites were summed within four prespecified classes: saturated fatty acids, unsaturated fatty acids, acylcarnitines and phosphatidylcholines/lysophosphatidylcholines for each mouse and analysed as log2(sum intensity + 1).

### Western blotting

Mouse liver tissues were lysed in protein lysis buffer (Beyotime, P0013B), and total protein concentrations were determined using a bicinchoninic acid (BCA) assay according to the manufacturer’s instructions. Protein samples were mixed with SDS–PAGE loading buffer (Beyotime, P0015L) and denatured at 100 °C for 3-5 min. Equal amounts of protein were separated by SDS-PAGE using 15% resolving gels with 5% stacking gels. Electrophoresis was performed at 85 V until the samples entered the resolving gel and subsequently at 130 V until adequate separation was achieved. Proteins were transferred onto methanol-activated polyvinylidene difluoride (PVDF) membranes (Millipore, IPVH00010) by wet transfer at a constant current of 300 mA for 60 min using transfer buffer containing 20% methanol. Membranes were blocked with 5% (w/v) non-fat milk in Tris-buffered saline containing Tween-20 (TBST) for 1 h at room temperature and subsequently incubated with primary antibodies against GAPDH (Proteintech, 60004-1-Ig), followed by incubation with the horseradish peroxidase (HRP) -conjugated goat anti-mouse IgG (Proteintech, SA00001-1) and chemiluminescent (Tanon, 180-5001) detection. After stripping and washing, the first membrane was reprobed with anti-PPARα (Santa Cruz Biotechnology, sc-398394). The second membrane was sequentially reprobed by anti-SCD1 (CST, 2438), anti-CYP4A14 (Proteintech, 11688-1-AP), ACSL1 (CST, 4047), and HMGCS2 (CST, 20940), with chemiluminescent detection performed after each appropriate secondary antibody incubation, anti-mouse or anti-rabbit IgG (Proteintech, SA00001-2). For each reprobing step, membranes were stripped using a stripping buffer (Beyotime, P0025), washed thoroughly with TBST, reblocked, and incubated with the next primary antibody. Immunoreactive bands were visualized using enhanced chemiluminescence reagent (Tanon, 180-5001) and imaged using a Tanon 4600 imaging system (Tanon). Band intensities were quantified using ImageJ. Protein abundance was normalized to GAPDH and expressed relative to the indicated control group.

### Statistical analysis and reproducibility

Statistical analyses were performed in R (v4.5.1) unless otherwise specified. The statistical methods, covariates, data transformations, and multiple-testing correction procedures used for individual analyses are described in the corresponding Methods sections. Unless otherwise stated, statistical tests were two-sided, and *P* < 0.05 was considered statistically significant. Where multiple hypotheses were tested, adjusted *P* values were reported as specified for the corresponding analysis. For animal experiments, each mouse was considered an independent biological replicate, and the numbers of animals included in individual analyses are indicated in the corresponding Methods sections and figure legends. For *in vitro* experiments, independent replicate cultures or reaction mixtures were analyzed as indicated for each experiment. Technical measurements were not treated as independent biological replicates.

## Data Availability

MetaCardis cohort: For the MetaCardis analyses in the present study, we used sequencing data retrieved from PRJEB37249 and restricted the analysis to healthy controls and UMMC participants that could be matched to the participant-level metadata reported in the original study, resulting in 270 healthy controls and 159 UMMC participants. Sample identifiers and phenotypic characteristics for the participants included in the present analyses are provided in Supplementary Table 1. The metabolomic data can be obtained from the raw article supplemental data.

Suzhou cohort: Metagenomic sequencing data for faecal samples can be accessed from the China Nucleotide Sequence Archive (CNSA) with the dataset identifier CNP0000175.

FBIP cohort: The raw metagenomic sequencing data generated in this study have been deposited in the Sequence Read Archive (SRA) database under accession code PRJNA1201249.

PPT and MED cohort: The raw microbiome samples generated in this study have been deposited in the European Nucleotide Archive (ENA) under accession code PRJEB64861.

The raw sequencing data generated from mouse liver tissues have been deposited in the NCBI Sequence Read Archive (SRA) under BioProject accession PRJNA1516159. The raw fecal shotgun metagenomic sequencing data from the mouse experiment have been deposited under BioProject accession PRJNA1517591. Annotated untargeted metabolomics data from mouse liver, plasma, and fecal samples are provided in Supplementary Table 5.

## Code Availability

All code used in this study can be accessed at https://github.com/jaspershen-lab/R.hominis_ohyA.

## Acknowledgments

This work was supported by start-up funding (Nanyang Assistant Professorship) provided to Dr. Xiaotao Shen from the Lee Kong Chian School of Medicine (LKCMedicine) and the School of Chemistry, Chemical Engineering, and Biotechnology (CCEB) at Nanyang Technological University, Singapore (NTU). This work was also supported by the Tier 1 grant from the Ministry of Education (MOE), Singapore, to Xiaotao Shen (Grant number: #025402-00001). Chuchu Wang was also supported by start-up funding (Nanyang Assistant Professorship) from LKCMedicine at NTU Singapore. Kazuyuki Kasahara was supported by the MOE Tier 1 (RG125/25) and Tier 2 (MOE-T2EP30223-0008) grants, and the National Medical Research Council (NMRC, MOH-001997). The authors declare the use of ChatGPT (OpenAI) for assistance with language polishing and grammar correction during the preparation of this manuscript.

## Author Contributions Statement

X.S., J.Z., and C.W. conceived the project. X.S. and C.W. supervised the study. J.Z. and C.Z. performed the public human cohort data analysis. C.H. and J.Z. performed the targeted LC-MS metabolomics data acquisition and analysis. L.H., M.H., and C.W. performed the protein-related experiments. S.T. and K.K. performed the bacterial cultivation experiments. J.Z., C.Z., X.S., and C.W. prepared all the figures. X.S., J.Z., C.Z., and C.W. wrote and revised the manuscript. All authors reviewed and contributed to the final manuscript.

