## Supplementary_information for "Gut *Roseburia hominis*-derived 10-hydroxystearic acid alleviates metabolic disorder by restoring hepatic fatty acid metabolism"

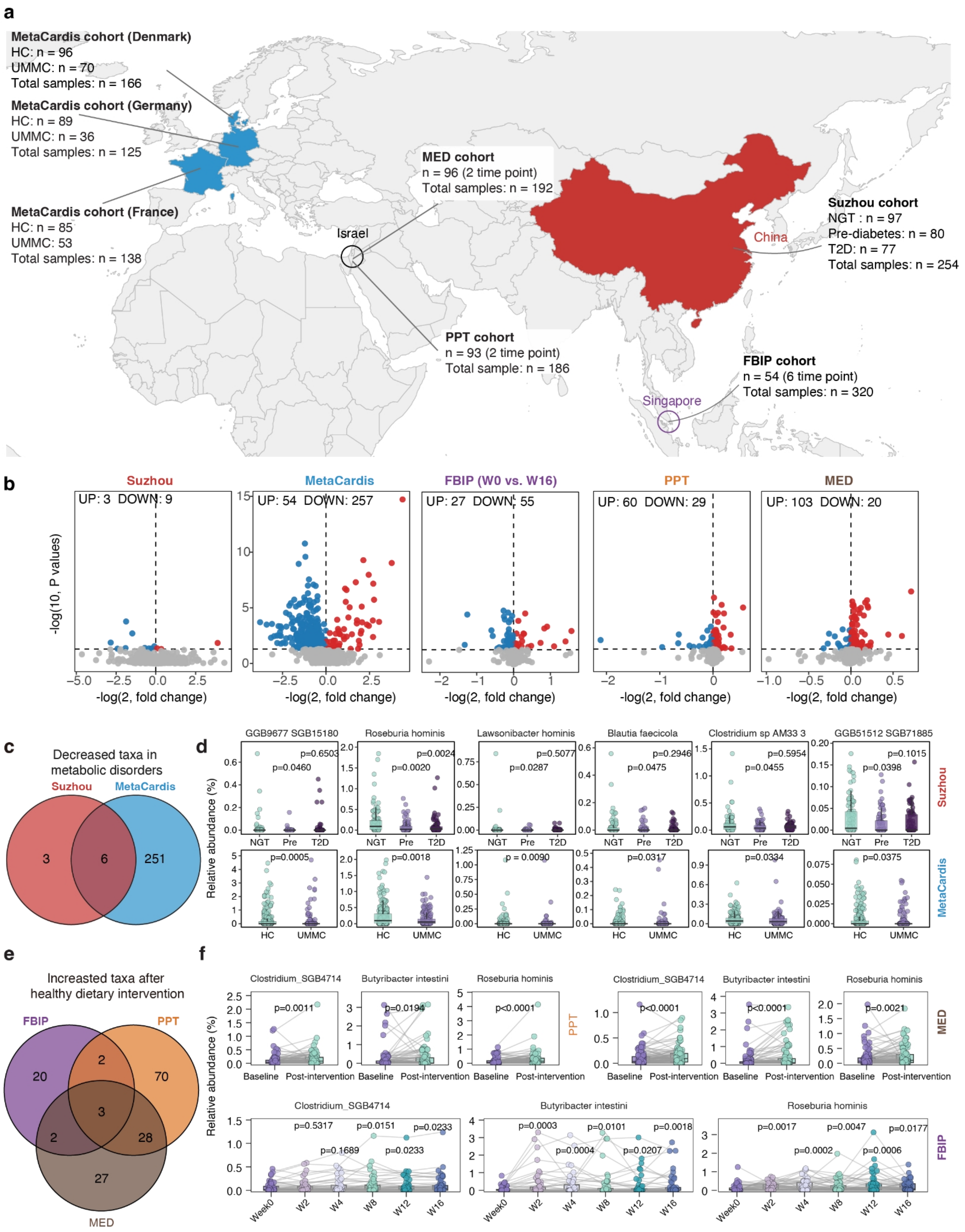


**Supplementary Figure 1 | Multi-cohort analysis identifies gut microbial taxa associated with metabolic disorders and dietary interventions.** (a) The distribution and source of the samples. (b) The volcano plots across 5 cohorts. (c) A Venn diagram showing gut taxa consistently decreased in metabolic disorder groups across the two metabolic disorder cohorts. (d) Relative abundances of representative metabolic disorder-associated taxa in the Suzhou cohort (top) and the MetaCardis cohort (bottom). (e) Venn diagram showing gut taxa consistently increased after dietary interventions across the FBIP, PPT, and MED cohorts. Only taxa exhibiting concordant post-intervention increases in all three cohorts were retained. (f) Relative abundances of representative diet-responsive taxa before and after dietary intervention in the FBIP, PPT, and MED cohorts.

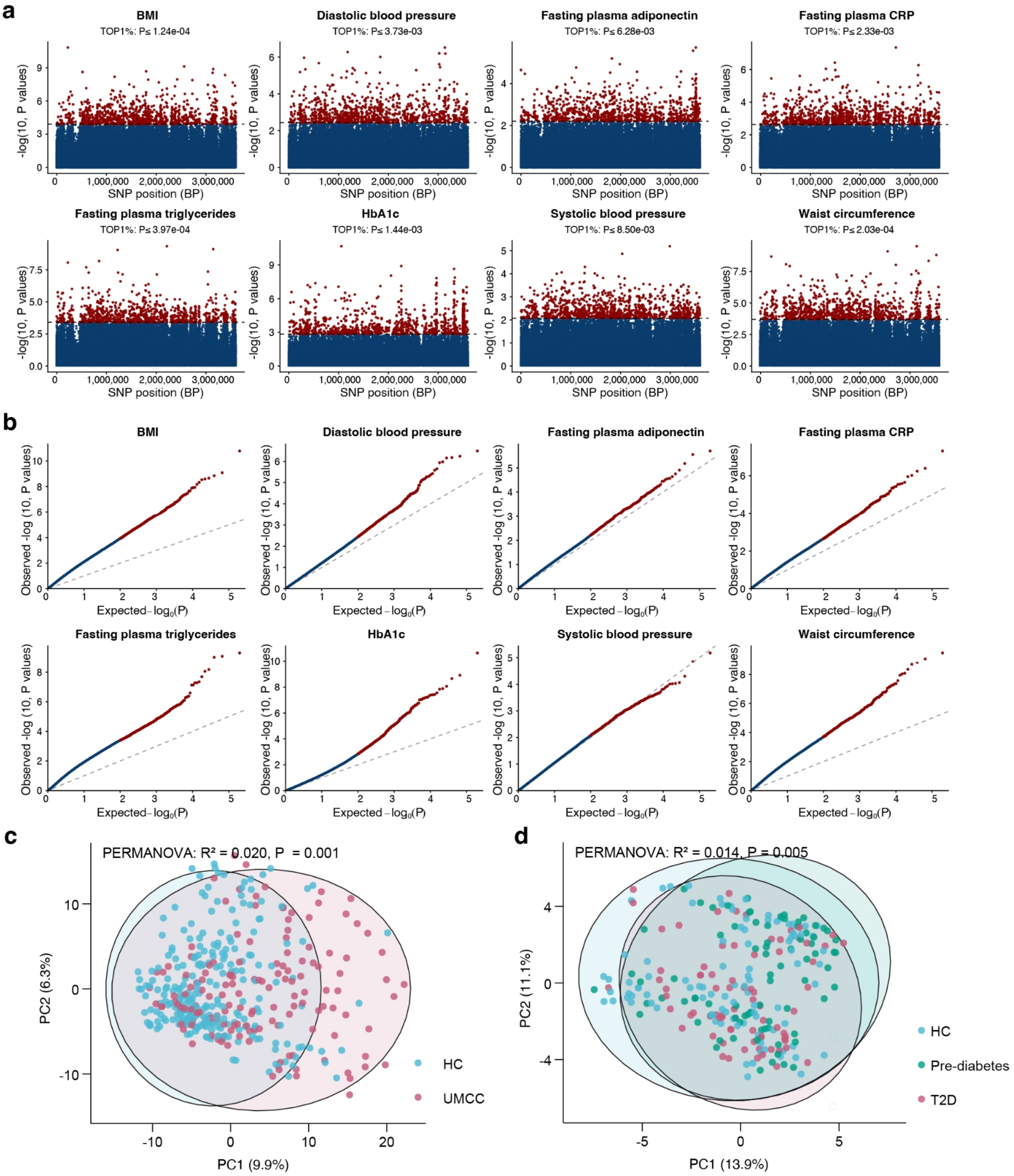


**Supplementary Figure 2 | Cardiometabolic trait-associated genetic variation in *R. hominis* distinguishes metabolic disorder status.** (a) Manhattan plots with top 1 % SNP *P* value cutoff showing the associations between SNPs and eight phenotypes: BMI, diastolic blood pressure, fasting plasma adiponectin, fasting plasma CRP, fasting plasma triglycerides, HbA1c, systolic blood pressure, and waist circumference. (b) Quantile-quantile (Q-Q) plots of the SNP association results for the eight phenotypes. (c) PCA of the MetaCardis cohort based on SNP profiles, comparing HC and UMMC individuals. (d) PCA of the Suzhou cohort based on SNP profiles, comparing NGT, Pre, and T2D individuals. Group differences in c and d were assessed by PERMANOVA with 999 permutations *P*-value.

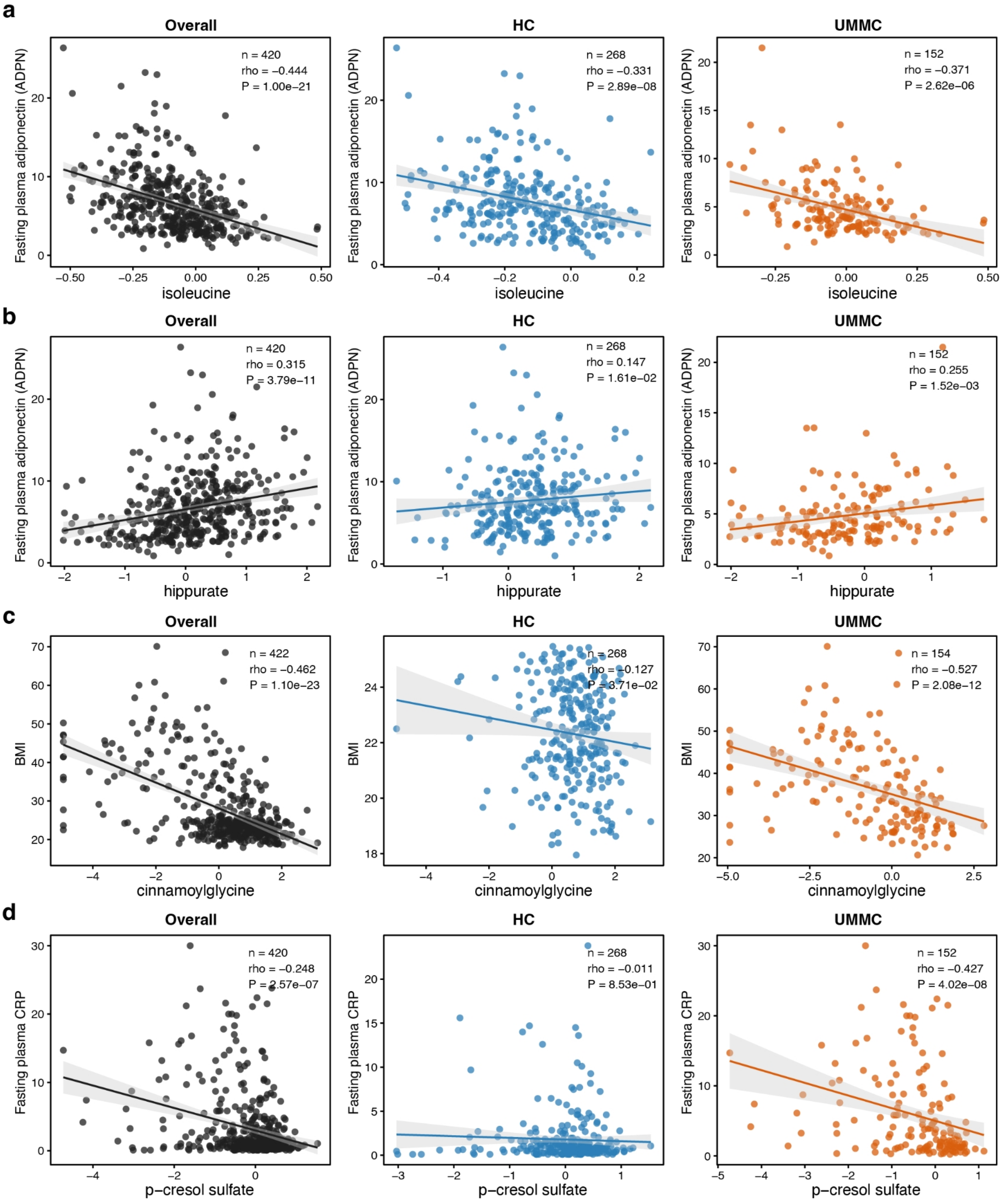


**Supplementary Figure 3 | Circulating metabolites associated with *ohyA* variants are linked to host cardiometabolic traits.** Scatter plots show the associations of (a) isoleucine with fasting plasma adiponectin, (b) hippurate with fasting plasma adiponectin, (c) cinnamoylglycine with BMI, and (d) *p*-cresol sulfate with fasting plasma CRP in the overall population, HC, and individuals with UMMC. Spearman's correlation coefficients and raw *P* values are shown. Lines indicate linear fits with 95% confidence intervals.

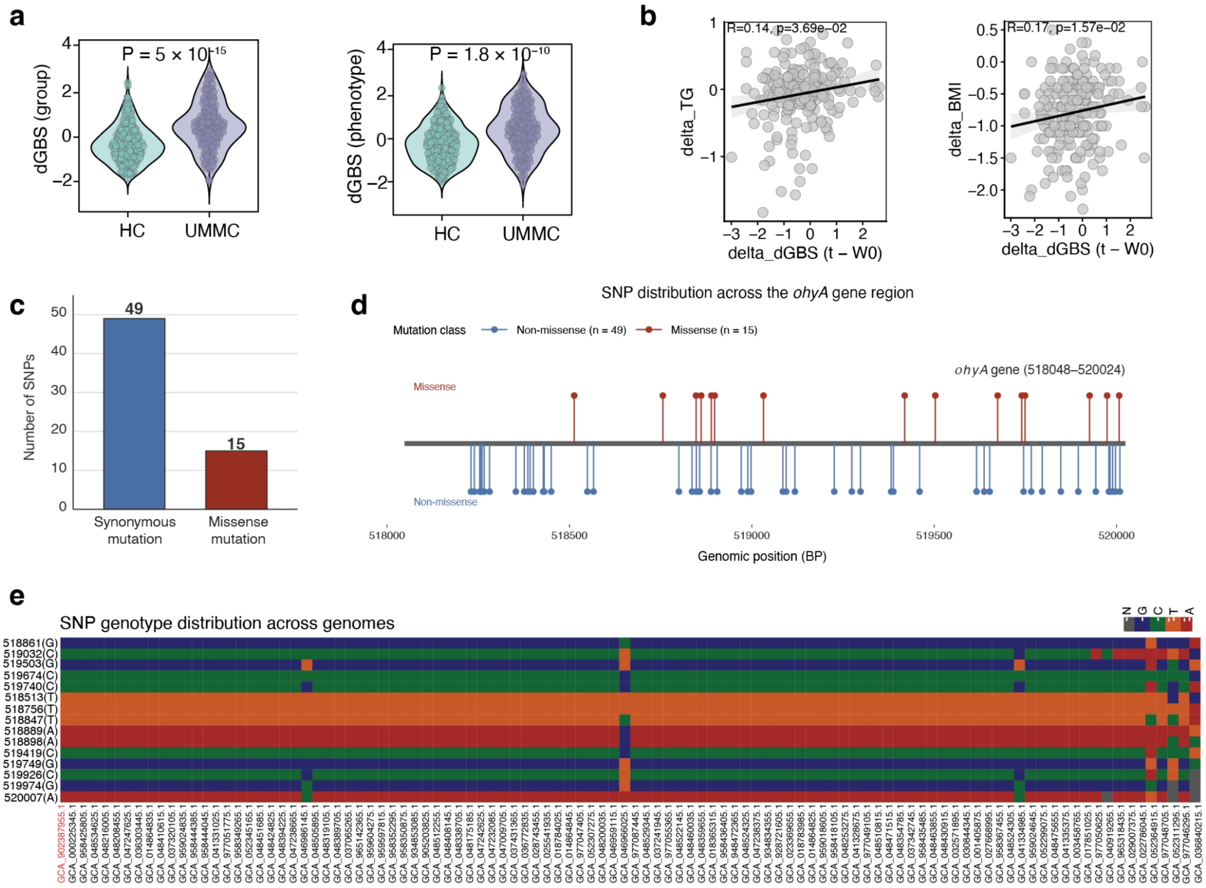


**Supplementary Figure 4 | *ohyA* genetic variation is associated with cardiometabolic status and dietary response.** (a) Comparison of group-derived and phenotype-derived *ohyA*-derived genetic burden scores (dGBS) between healthy controls (HC) and individuals with UMMC in the MetaCardis cohort. (b) Associations between within-participant changes in phenotype-derived dGBS (ΔdGBS; time point − baseline [W0]) and corresponding changes in BMI (ΔBMI) or triglycerides (ΔTG) after dietary intervention in the FBIP cohort. (c) The mutation type of 64 SNPs in *ohyA*. (d) The 64 SNP positions in *ohyA*. (e) Distribution of the 15 *ohyA* missense variants identified in the study across available *R. hominis* RefSeq genomes.

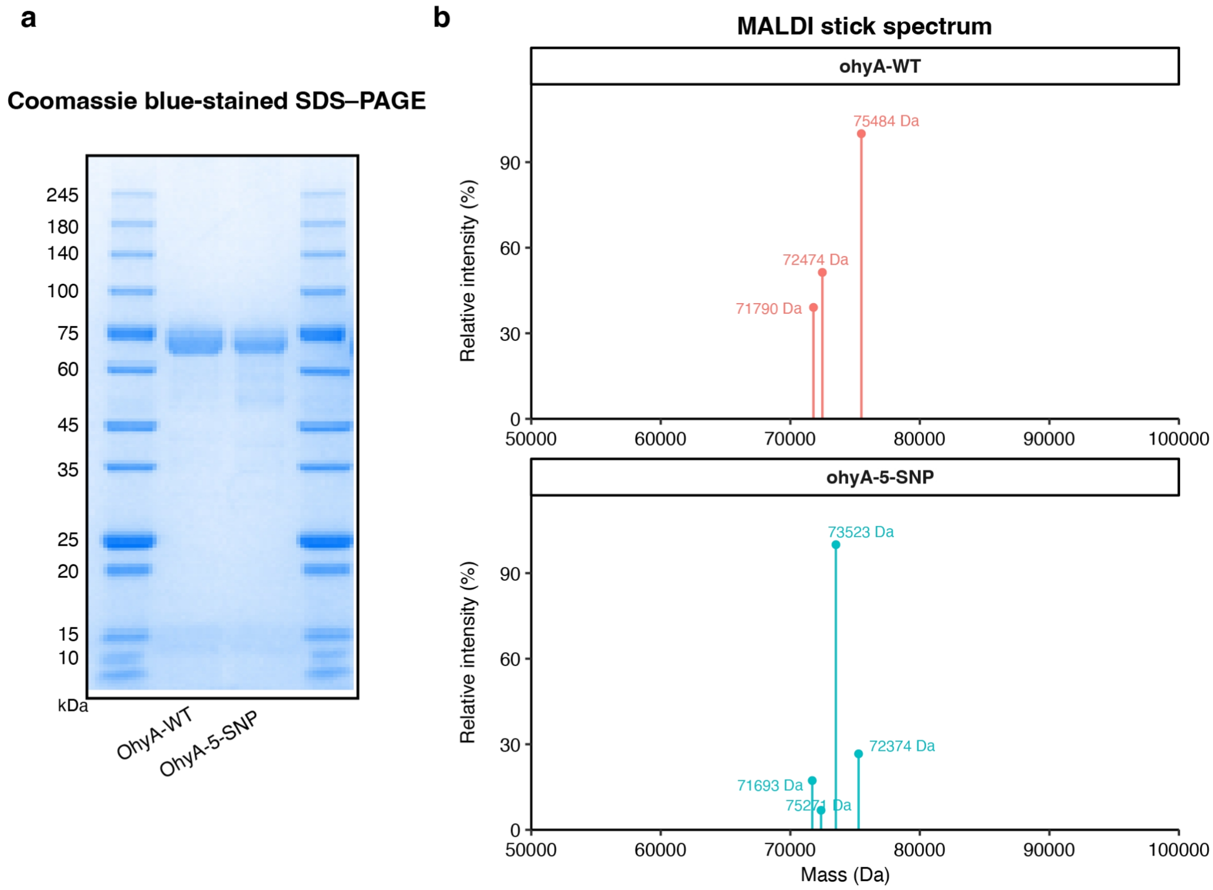


**Supplementary Figure 5 | Biochemical characterization of recombinant wild-type and variant OhyA proteins.** (a) Coomassie blue-stained SDS–PAGE of purified WT and 5-SNP OhyA proteins. (b) MALDI mass spectra of the purified WT and 5-SNP OhyA proteins, with major molecular-mass peaks indicated.

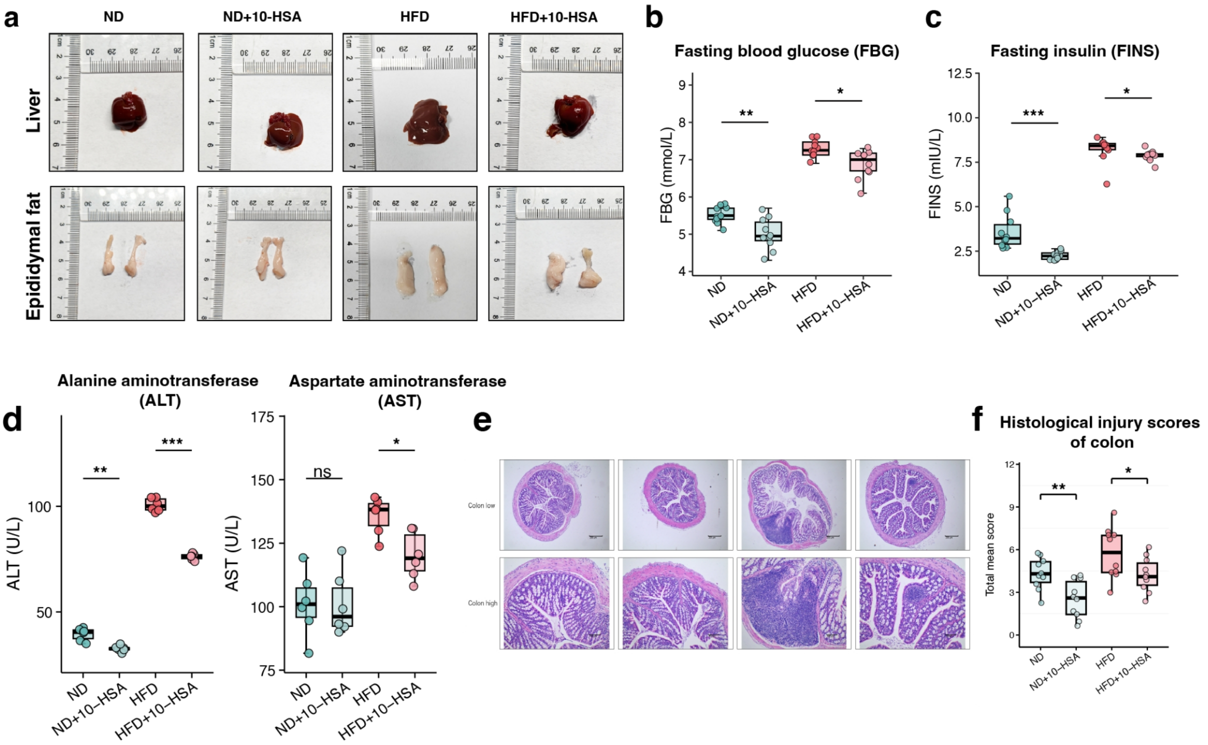


**Supplementary Figure 6 | 10-HSA improves metabolic and hepatic outcomes in HFD-fed mice.** (a) Representative images of liver and epididymal adipose tissue. (b-c) Fasting blood glucose (FBG) and fasting insulin (FINS) levels. (d) Serum alanine aminotransferase (ALT) and aspartate aminotransferase (AST) levels as indicators of liver function. (e) Representative H&E-stained colon sections. (f) Histological injury scores of the colon.

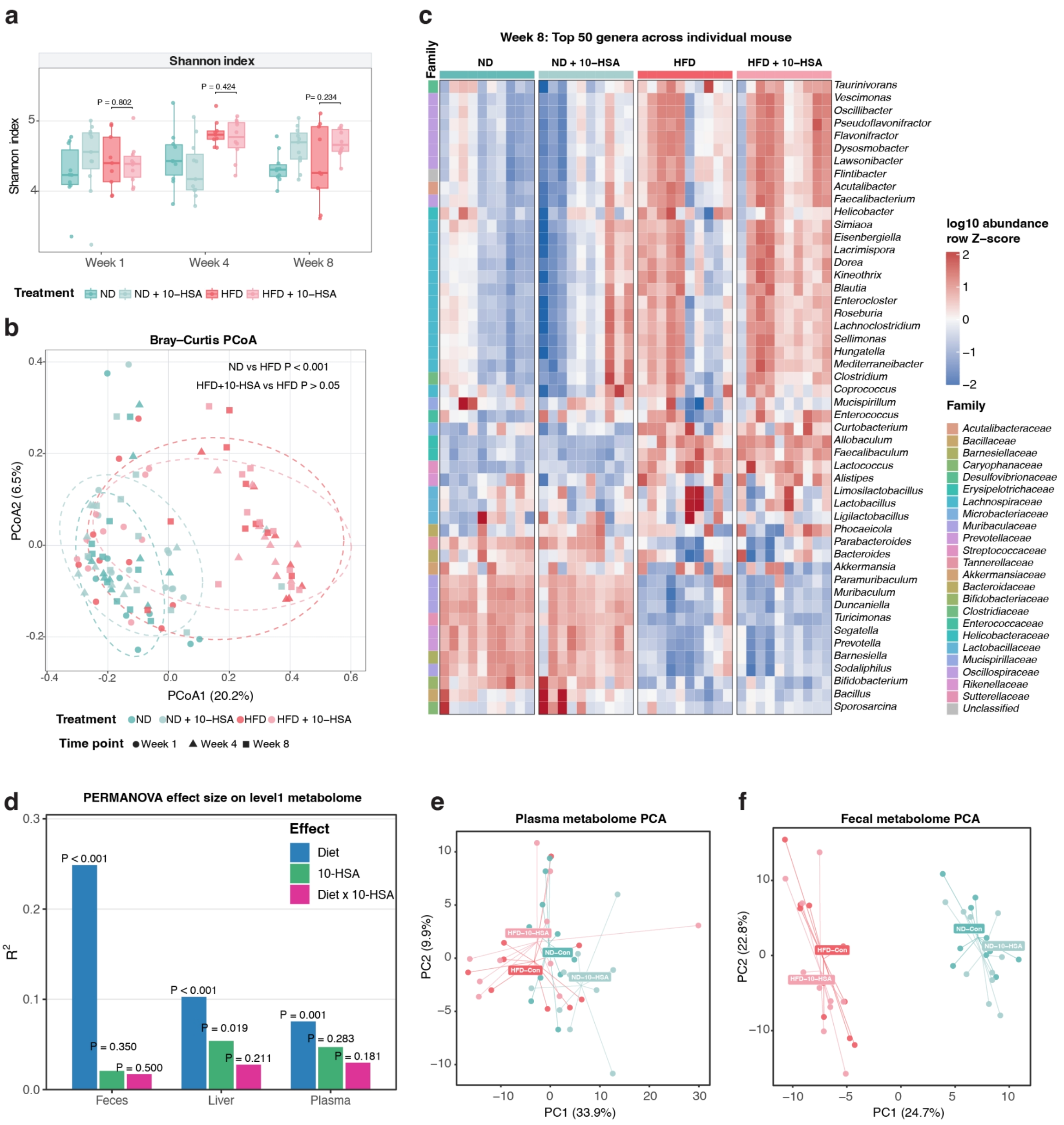


**Supplementary Figure 7 | 10-HSA exerts limited effects on the gut microbiota but induces tissue-specific metabolic responses.** (a) Shannon diversity of the gut microbiota at weeks 1, 4, and 8. (b) Bray-Curtis PCoA of gut microbial composition across treatment groups and time points. (c) Heatmap showing the relative abundance of the top 50 bacterial genera at week 8. (d) PERMANOVA effect sizes of diet, 10-HSA, and their interaction on fecal, liver, and plasma metabolomes. PCA of plasma (**e**) and fecal (**f**) metabolomic profiles across the four treatment groups.

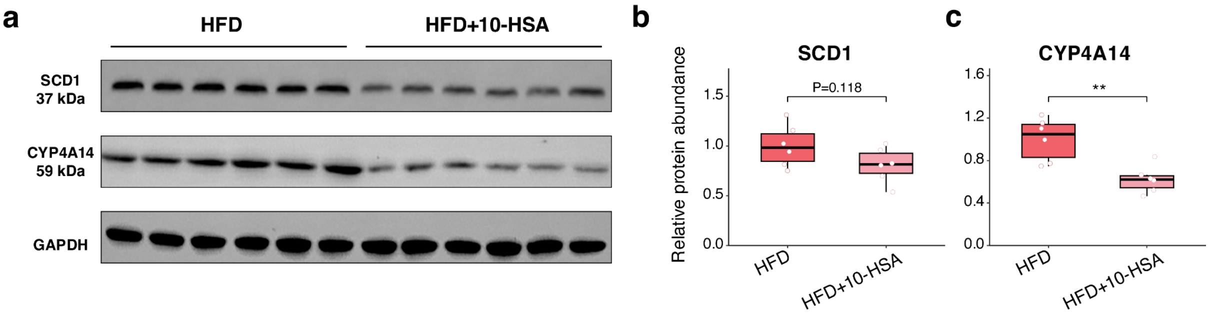


**Supplementary Figure 8 | 10-HSA supplementation decreases hepatic SCD1 and CYP4A14 protein abundance despite transcriptional upregulation.** (a) Representative western blot images of SCD1 and CYP4A14 in liver tissues from HFD and HFD + 10-HSA mice, with GAPDH shown as the loading control. Densitometric quantification of hepatic SCD1 (b) and CYP4A14 (c) protein abundance normalized to GAPDH. Each dot represents an individual mouse; bars indicate the mean ± s.e.m. $n = 6$ mice per group.
